# Human-specific features of the cerebellum and ZP2-regulated synapse development

**DOI:** 10.1101/2025.09.08.674970

**Authors:** Suel-Kee Kim, Adriana Cherskov, Aastha Sindhwani, Sang-Hun Choi, Hyojin Kim, Ming-Li Li, Menglei Zhang, Xoel Mato-Blanco, Yuting Liu, Nicola Micali, David M. Young, Mark Estacion, Yueqi Zhang, José Manuel Ruiz-Jiménez, Anandita Nadkarni, Victor Luria, Suvimal Kumar Sindhu, Ipsita Chatterjee, Akemi Shibata, Dan Liang, Hyesun Cho, Saejeong Park, Ana Spajic, Rothem Kovner, Martina Glavan, Rachel J. Chen, Ryan D. Risgaard, Xinyun Li, Sirisha Pochareddy, Amir Karger, Anita Huttner, Yury M. Morozov, Etienne W. Daadi, Carlo Colantuoni, Kevin T. Gobeske, John J. Ely, Patrick R. Hof, Marcel M. Daadi, Chet C. Sherwood, Alvaro Duque, Shaojie Ma, Andre M. M. Sousa, Stephen G. Waxman, Pasko Rakic, Gabriel Santpere, Stephan J. Sanders, Nenad Sestan

**Affiliations:** Department of Neuroscience, Yale School of Medicine, New Haven, CT, USA; Hospital del Mar Research Institute, Parc de Biomèdica de Barcelona (PRBB), 08003 Barcelona, Catalonia, Spain; Department of Psychiatry and Behavioral Sciences, UCSF Weill Institute for Neurosciences, University of California, San Francisco, CA, USA; Department of Neurology and Center for Neuroscience and Regeneration Research, Yale School of Medicine, New Haven, CT, USA; Rehabilitation Research Center, Veterans Affairs Connecticut Healthcare System, West Haven, CT, USA; Waisman Center, University of Wisconsin-Madison, Madison, WI, USA; IT-Research Computing, Harvard Medical School, Boston, MA, USA; Southwest National Primate Research Center, Texas Biomedical Research Institute, Departments of Cell Systems and Anatomy, and Radiology, Long School of Medicine, University of Texas Health at San Antonio, San Antonio, TX, USA; Departments of Neurology and of Neuroscience, Johns Hopkins School of Medicine, Baltimore, MD, USA; Institute for Genome Sciences, University of Maryland School of Medicine, Baltimore, MD, USA; MAEBIOS Epidemiology Unit, Alamogordo, NM, USA; Department of Anthropology and Center for the Advanced Study of Human Paleobiology, The George Washington University, Washington, DC, USA; Nash Family Department of Neuroscience, Friedman Brain Institute and Center for Discovery and Innovation, Icahn School of Medicine at Mount Sinai, New York, NY, USA; Institute of Neuroscience, CAS Center for Excellence in Brain Science and Intelligence Technology, University of Chinese Academy of Sciences, Shanghai, China; Department of Neuroscience, University of Wisconsin-Madison, Madison, WI, USA; Institute of Developmental and Regenerative Medicine, Department of Paediatrics, University of Oxford, Oxford, UK; New York Genome Center, New York, NY, USA; Department of Comparative Medicine, Department of Genetics, Department of Psychiatry, Wu Tsai Institute, Program in Cellular Neuroscience, Neurodegeneration and Repair, and Yale Child Study Center, Yale University, New Haven, CT, USA

**Keywords:** Brain evolution, Cerebellum, Human-specific, Synapse development, ZP2, Multiomics

## Abstract

Understanding the unique features of the human brain compared to non-human primates has long intrigued humankind. The cerebellum refines motor coordination and cognitive functions, contributing to the evolutionary development of human adaptability and dexterity. To identify shared and divergent features across primates, we conducted single-nucleus transcriptomic and chromatin accessibility profiling of the adult cerebellar cortex in humans, chimpanzees, macaques, and marmosets. We revealed human-specific transcriptomic and regulatory features, particularly those involved in synaptogenesis. Notably, we identified an enrichment of the sperm receptor zona pellucida glycoprotein 2 (ZP2) and its potential interactors, known for their roles in gamete interaction, in human granule cells. Experimental data show that ZP2 expression in human granule cells is induced by pontine mossy fibers, reducing synaptic proteins at pontocerebellar glomerular synapses, and decreasing cerebellar neuron electrophysiological activity. This unexpected co-option of ZP2 in human-specific synapse regulation provides insights into the evolutionary specialization of the human cerebellum.

## INTRODUCTION

The cerebellum plays a critical role in controlling motor function, attention, and cognitive processes, with its outputs targeting motor and cognitive regions of the thalamus and cerebral cortex^1–4^. Cortical excitatory projection neurons regulate cerebellar activity via glutamatergic excitatory pontine neurons, which form mossy fibers that synapse with excitatory granule and inhibitory Golgi neurons in the cerebellar cortex^5^. Abnormal development or dysfunction of this circuitry is linked to motor impairments, such as ataxia and complex neuropsychiatric conditions, including attention-deficit hyperactivity disorder (ADHD), autism spectrum disorder (ASD), schizophrenia (SCZ), and general liability for cognitive disorders^2,3,6–8^.

Recent advancements in next-generation sequencing have facilitated detailed investigations of cell type-specific molecular processes underlying evolutionary specialization in the primate and human nervous systems. While single-cell analyses have begun to uncover genomic profiles of the cerebellum in various mammals^9–17^, no study has integrated multiple modalities to compare humans with chimpanzees, our closest living relatives, alongside other non-human primates. This study overcomes these limitations by generating the first single-nucleus multiomic (paired snRNA-seq and snATAC-seq) atlas of the adult cerebellar cortex encompassing four major anthropoid phylogenetic groups: humans (*Homo sapiens*), chimpanzees (*Pan troglodytes*), rhesus macaques (*Macaca mulatta*), and common marmosets (*Callithrix jacchus*) (https://nemoanalytics.org/p?l=SestanPrimateCerebellum). This resource enables a robust examination of both convergence and divergence in gene expression and chromatin accessibility at the single-cell resolution. Incorporating chimpanzees and two outgroup species allows reliable identification of cell type-specific features shared across primates, specific to certain primate groups or species, or unique to humans.

## RESULTS

### Single-cell Multiomic Atlas of the Anthropoid Cerebellar Cortex

We conducted paired snRNA-seq and snATAC-seq on histologically verified postmortem posterolateral cerebellar cortex (Crus I and II/lobule VI and VII) from neurotypical humans (n=4), chimpanzees (n=3), rhesus macaques (n=4), and common marmosets (n=4) (Figure 1A, Table S1). After stringent quality control, we obtained 69,302 nuclei with high-quality RNA profiles, detecting an average of 2,232 genes per nucleus and 63,491 nuclei in the ATAC assay, with 60,824 nuclei overlapping between both assays (Figure S1A and S1B).

**Figure 1.**
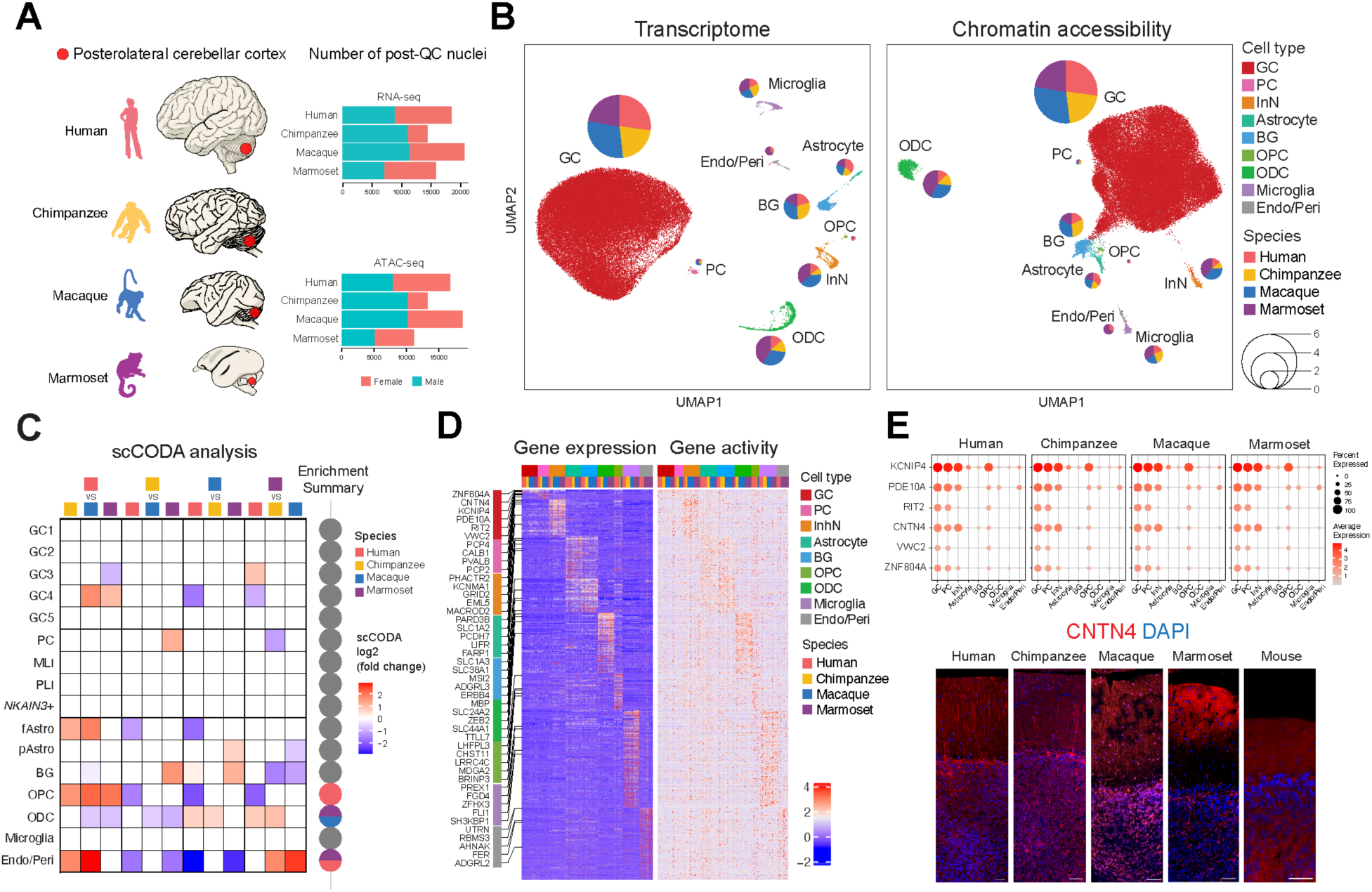
Single-nucleus multiome analysis of primate cerebellar cortex. (A) Overview of primates included in the study and the number of post-QC nuclei obtained for multiome data analysis. (B) UMAP plots for Multiome data. Color codes and cell-type abbreviations are consistently applied throughout all figures. GC: granule cells, PC: Purkinje cells, InN: inhibitory interneurons, BG: Bergmann glial cells, Astrocyte, OPC: oligodendrocyte precursor cells, ODC: oligodendrocytes, Microglia, Endo/Peri: endothelial cells and pericytes. Pie charts illustrate the species composition within each cell type, with the size of each pie reflecting the proportion of that cell type among all cells. (C) Cell type proportions. Left: Heatmap illustrating scCODA-corrected fold changes in subtype abundance between species pairs. Non-zero values represent significant changes. Right: Pie charts summarizing species-specific enrichment patterns of subtype abundance. (D) Heatmap showing cell type–specific gene expression conserved across species, reflecting shared molecular features of core cell identity. The top five marker genes for each cell type are labeled. (E) Expression of conserved genes in GCs. Top: Expression of the six GC-enriched genes exhibiting high expression levels and conserved patterns across primates. Bottom: Localization of CNTN4 protein in adult cerebellum across four primates and mice. Scale bar, 50 μm. See also Figures S1-3.

To establish a cellular taxonomy of the cerebellum across primates, we integrated transcriptome data using unsupervised clustering and Uniform Manifold Approximation and Projection (UMAP). Leveraging canonical marker expression profiles^9,13–15,17^, we identified nine major clusters representing putative cell classes (Figure 1B). These clusters showed consistent separation and neighborship across both snRNA- and snATAC-defined UMAP dimensions and species-independent integration (Figure S1C). All major cell classes were detected in all four primates and validated by canonical cell type-specific gene expression and corresponding chromatin accessibility patterns (Figures S1D and S1E). Neuronal clusters represented granule cells (GCs, *NEUROD1*/*GABRA6/SLC17A7*), Purkinje cells (PCs, *CALB1/ALDOC*), and inhibitory interneurons (InNs, *GAD1*/*GAD2)*. Non-neuronal clusters included astrocytes (*GFAP*/*SORCS2*), Bergmann glial cells (BGs, *SLC1A3*), oligodendrocyte precursor cells (OPCs, *PDGFRA*), oligodendrocytes (ODCs, *MOBP*/*MBP*), microglia (*P2RY12*), and endothelial cells and pericytes (Endo/Peri, *PECAM1*). The classification accuracy was further validated by comparing global transcriptomic profiles with published scRNA-seq datasets of adult mouse and human cerebellum^15,17^ (Figures S1F and S1G).

Further transcriptomic subclustering revealed refined subtypes within major cell classes, reflecting potential heterogeneity in spatial distribution, functionality, and physiological states. Astrocytes were divided into protoplasmic and fibrous astrocyte subtypes, while inhibitory neurons included PCs, molecular layer interneurons (MLI), Purkinje layer interneurons (PLI), and an unknown subtype marked by high *NKAIN3* expression (Figures S2A and S2B). GCs were divided into five subclusters, showing high transcriptomic similarity without distinct expression of cerebellar microzone markers such as *ALDOC* or *NOS1*^18^ (Figures S2C and S2D).

The proportion of cell types and subclusters was generally consistent across donors and species (Figures 1B, 1C, S1A, S1B, and S2E), with the exception of OPCs, ODCs, and Endo/Peri clusters. Humans showed a higher proportion of OPCs and a lower proportion of ODCs than other primates (Figure S2E). Our dataset, restricted to adult donors (ages 36 to >50 years), revealed no clear age-related trend and does not capture earlier developmental progression (Figure S2F). However, previous studies have reported delayed oligodendrocyte maturation and myelination, and OPCs have been shown to persist into adulthood in the human cerebral cortex^16,19–21^. Our findings add to these observations by showing a higher proportion of OPCs in the human cerebellum relative to other primates, consistent with prolonged OPC persistence in human adults. Given the relatively low abundance of these cell types in the adult cerebellum, cautious interpretation is warranted. In contrast, GCs, the most abundant neuronal cell type in the brain, constitute up to 99% of all neurons in our primate cerebellum dataset. Previous studies have reported that the GC-to-PC ratio increases in species with larger cerebella^4,18^. Given this composition, the low proportions of other cell types (5.5% glia and 1.5% InNs) were expected. These findings confirm that all major cerebellar cell types are conserved across the four anthropoid primates.

### Conserved Cell Type-Specific Molecular Features

We next assessed conserved molecular features across primates at cell type-specific resolution, focusing on genes that define core cell identity (Figures 1D, S3A, and S3B, Table S1). Conserved cell type-specific genes were defined as those showing enrichment in the same cell type across all four species. This approach allowed us to focus on genes related to cell type identity that are evolutionarily stable. Consistent with these criteria, the vast majority of conserved genes were restricted to a single cell type. Microglia showed the highest number of genes with conserved cell type-specific expression, followed by ODCs and Endo/Peri cells, while GCs and PCs displayed few genes (Figure S3A). Given the predominance of GCs in the adult cerebellum, we focused on their conserved features to identify fundamental characteristics shared across the primate cerebellum. Top GC-enriched conserved genes included *KCNIP4, PDE10A, RIT2, CNTN4, VWC2,* and *ZNF804A* (Figures 1D, 1E and S3B). Notably, *KCNIP4, CNTN4,* and *ZNF804A* have been associated with synaptic functioning and neuropsychiatric disorders (https://omim.org, https://gene.sfari.org), suggesting their potential involvement in cerebellar function. *KCNIP4* has also been associated with cerebellar ataxia in dogs^22^. Chromatin accessibility profiles revealed conserved chromatin-accessible peaks near the promoters of these genes across primates (Figure S3C). Notably, *VWC2*, previously reported for its higher expression in humans compared to mice^23^, showed primate-specific conserved expression in GCs, while CNTN4 was expressed in all four primates studied and the marsupial opossum but was absent in mice, suggesting its evolutionary significance (Figures 1E, S3B, and S3D). While this study primarily focuses on defining human-specific molecular features, our single-cell data from multiple primate species provides a valuable resource for uncovering both conserved and unique characteristics of the primate cerebellum.

### Divergent Cell Type-Specific Molecular Features

We next analyzed species-specific transcriptomic changes, identifying 744 genes upregulated and 249 downregulated in humans, along with 611 upregulated and 312 downregulated genes in Hominini, which were shared between humans and chimpanzees but distinct from the other two primates (Figures 2A, S4A, Table S1). In microglia, human-specific upregulation included neuropsychiatric disorder-associated genes *FOXP2, DSCAM,* and *CACNA1D*, aligning with our previous findings of their human-specific expression in the prefrontal cortex (PFC)^24^. Most human- or Hominini-specific expressions were enriched in a single cell type, with GCs displaying the highest number, possibly due to their abundance and heterogeneity compared to other cell types. To further quantify overall transcriptomic differences between species, we computed cross-species transcriptomic divergence within each cell type. To minimize potential biases related to differences in cluster size or cell abundance, we excluded underrepresented PC, OPC, and Endo/Peri clusters and randomly selected equal numbers of cells from the remaining cell types. Notably, GCs and microglia exhibited the most pronounced transcriptomic divergences, with marmosets showing the greatest distance from humans (Figure 2B). These results indicate that in GCs, this divergence is largely driven by a high number of species-specific genes (Figure S4A). In microglia, however, the divergence is explained by a smaller number of genes that undergo large changes in expression. Although the cerebellum is recognized as one of the most evolutionarily conserved brain regions^25^, our findings suggest that GCs may have been subject to higher evolutionary pressures in humans.

**Figure 2.**
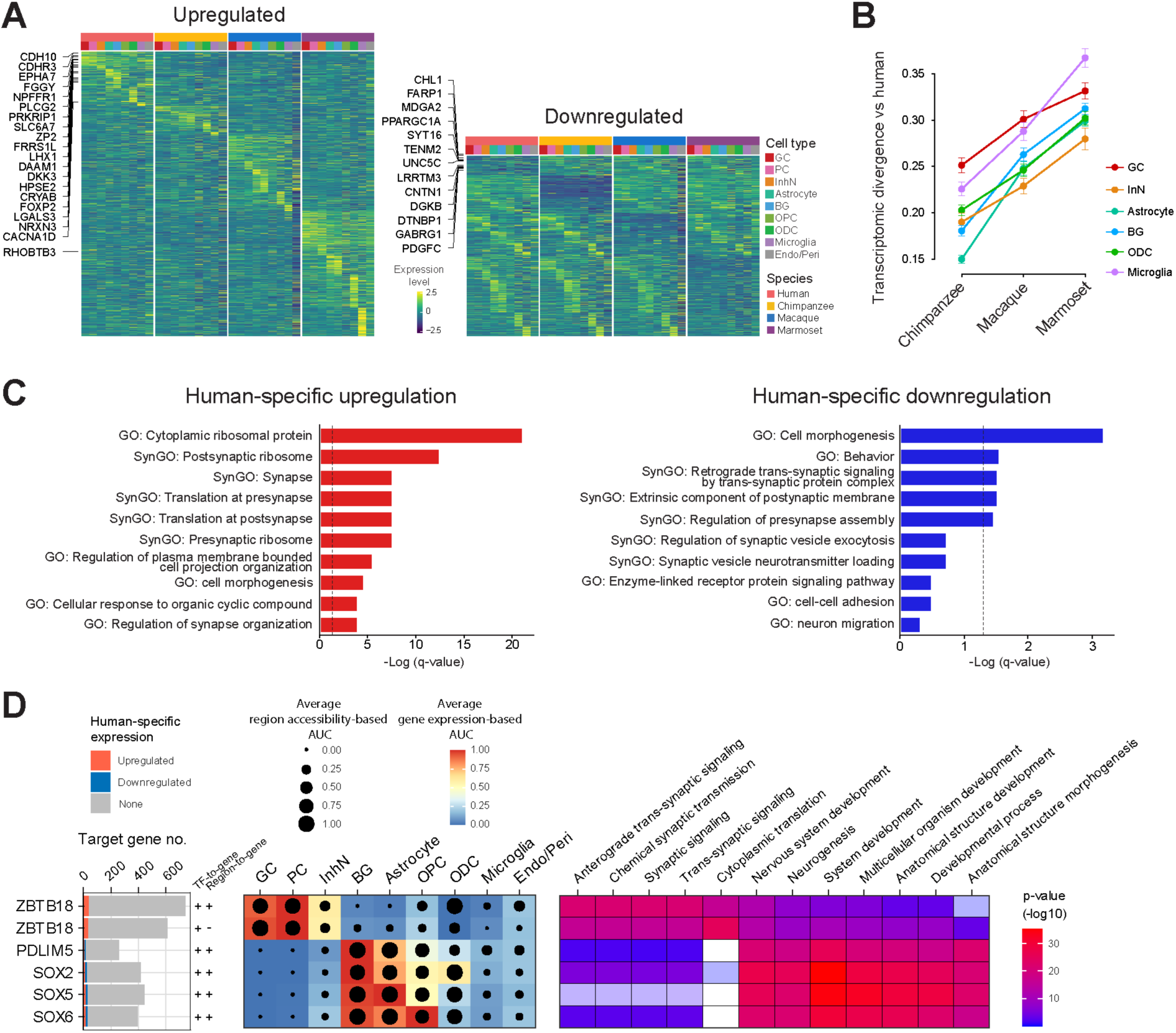
Human-specific and species-divergent molecular features. (A) Heatmap showing species-divergent gene expression across cell types, illustrating species-biased expression patterns within each cell type. Genes with human-specific upregulation or downregulation in GCs are labeled. (B) Transcriptomic divergence of each species compared to human within each cell type. (C) Enriched gene ontology (GO) and Synaptic GO (SynGO) terms for genes with human-specific expression. Vertical dotted lines indicate a significance of p = 0.05. (D) Predicted transcription factor (TF) regulatory networks regulating human-specific expression and GO enrichment analysis for each regulon. “+” and “–“ indicate TF-to-gene and region-to-gene correlation coefficients, where a positive correlation for both suggests that TFs open chromatin and activate gene expression. See also Figures S4 and S5.

Gene ontology (GO) and Synaptic GO (SynGO) analyses of genes with species-divergent expression revealed that upregulated genes in humans and Hominini were enriched for terms related to synaptic protein translation and localization, cell projection and synapse organization, and cytoskeleton regulation, particularly in GCs (Figures 2C, S4B and S4C, Table S2). Many ribosomal protein genes, especially those expressed at synaptic membranes and involved in local translation at the synapse^26^, displayed human- and Hominini-specific upregulation in GCs, InNs, and BGs. Human- and Hominini-specific downregulated genes were also enriched for terms associated with synapse formation and function, including trans-synaptic protein complex, regulation of presynapse assembly and synapse vesicle exocytosis, and cell-cell adhesion. These findings suggest potential roles of the human- and Hominini-specific gene expressions in synaptogenesis and synaptic function.

To investigate whether convergent upstream mechanisms drive human-specific and cell type-specific expressions, we utilized the SCENIC+ approach to construct the transcription factor (TF) regulatory network by integrating gene expression and chromatin accessibility profiles of the human cerebellum^27^. This analysis identified putative regulons, representing groups of target genes regulated by core TFs across distinct cerebellar cell types (Figure S5A). Among these identified regulons, five were significantly enriched with target genes exhibiting human-specific expression (Figure 2D). Notably, *ZBTB18* emerged as a core TF in GCs, PCs, and InNs, regulating over 600 target genes, which significantly intersected with human-upregulated genes. GO analyses indicated that *ZBTB18* targets are highly enriched in synaptic transmission and signaling pathways (Figure 2D), aligning with previous findings that highlight its critical role in human synaptic organization and neuronal connectivity^28,29^. Given the high enrichment of synapse-related terms among genes with human-specific regulation in GCs (Figure S4B), we further investigated their potential regulation by *ZBTB18*. Of the 31 genes with human-specific expression and synaptic function in GCs, nine were identified as putative *ZBTB18* targets, including three ribosomal proteins involved in local translation at the synapse (Figure S5B). In addition, *PDLIM5*, *SOX2*, *SOX5*, and *SOX6* were identified as core TFs in glial cells, each targeting a significant number of genes with human-specific expression (Figure 2D). These results highlight multiple key regulatory networks that may drive human-specific gene expression in the cerebellum.

### Human-Specific Expression of ZP2 and Oocyte-Sperm Interaction Genes

Having identified genes with human-specific expression (Figure 2B, Table S1), we next examined which of these genes also showed cerebellar-specific expression across brain regions. Particularly intriguing among the genes differentially expressed in GCs was *zona pellucida glycoprotein 2* (*ZP2*), known as a sperm receptor^30,31^. *ZP2* showed the strongest cerebellar specificity among the human-enriched genes in GCs, with expression markedly higher in the cerebellum than in other brain regions (Figure 3A). Its potential function in the brain has not been previously investigated, making it an especially compelling candidate for further analysis. Notably, our analysis of multiple published datasets confirmed *ZP2* expression in the human cerebellum, with surprisingly higher levels compared to the ovary^16,25,32–36^ (Figures 3B and S6A). In line with our previous findings^25^, *ZP2* transcripts were detected in our snRNA-seq dataset, and abundant protein presence was observed in tissue sections using various ZP2 antibodies, predominantly within human GCs (Figures 3C and 3D). Gene expression appeared less pronounced when assessed via in situ hybridization (Figure S6B) and was detected in less than 20% of GC populations in our snRNA-seq dataset (Figure 3C). This discrepancy between gene and protein levels could be attributed to snRNA-seq dropout events and the notably extended half-life of ZP proteins^37^.

**Figure 3.**
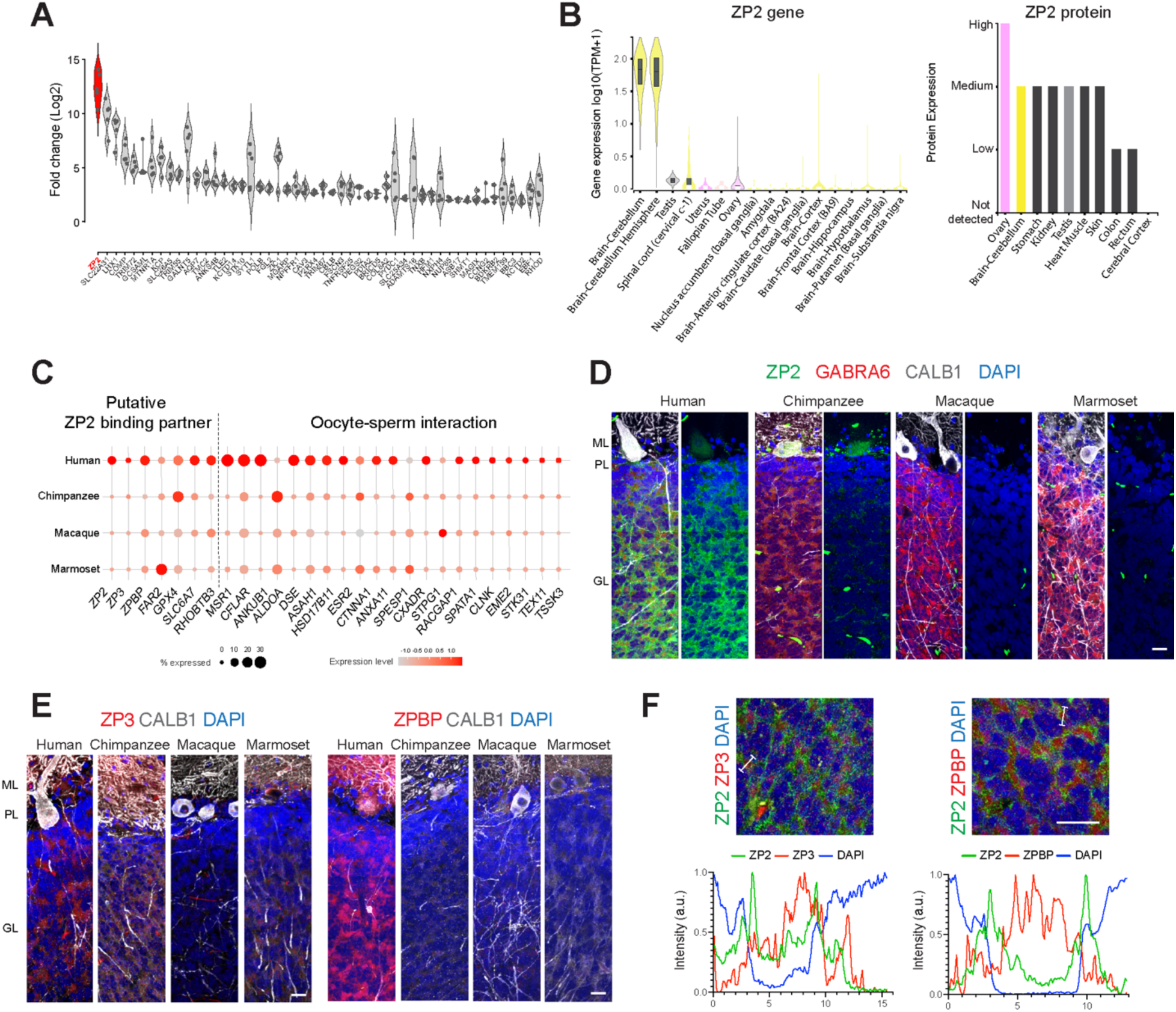
Human-specific expression of ZP2 and oocyte-sperm interaction genes. (A) Fold-change expression analysis of the top 50 human-specific upregulated genes, comparing the cerebellum to other brain regions, highlighting cerebellum-specific upregulation of *ZP2*. Brain regions analyzed include the neocortex (NCX), mediodorsal nucleus of the thalamus (MD), hippocampus (HIP), striatum (STR), and amygdala (AMY)^16,48^. (B) Left: *ZP2* gene expression across various brain regions and organs in adult humans^33^. Right: ZP2 protein expression levels across organs (https://www.proteinatlas.org/). (C) Expression of putative ZP2 binding partner and oocyte-sperm interaction genes in GCs, showing human- and Hominini-specific expression. (D) Immunohistochemical analysis of ZP2, GC marker GABRA6, and PC marker CALB1 in adult primate cerebellar cortex. (E) Localization of ZP3 and ZPBP proteins across four primates. (F) Localization of ZP2, ZP3, and ZPBP proteins within the cerebellar glomeruli. The spatial distribution of their signal intensity along the marked line is displayed at the bottom. Scale bars 20 µm. See also Figure S6.

The role of ZP2 has been extensively studied in mammalian reproduction, where it acts as a crucial regulator for oocyte-sperm binding, fertilization, and prevention of polyspermy^30,31^. ZP2 forms long polymers with other ZP family members, ZP1-4, to constitute the zona pellucida, the extracellular matrix (ECM) enveloping the egg, and interacts with ZPBP on acrosome-reacted sperm^30^. Consequently, we were intrigued to explore whether these and other putative ZP2 binding partners, predicted from protein interaction databases (BioGRID, IntAct, STRING, and FpClass), are expressed in the cerebellum and/or exhibited human-specific expression. We observed that several genes encoding validated or predicted ZP2 interactors are indeed expressed in the cerebellum (Figures S6C-E). Interestingly, among these, *ZP3*, *ZPBP*, *SLC6A7,* and *RHOBTB3* displayed human-specific upregulation, and *GPX4* showed Hominini-specific upregulation in GCs (Figures 3C, S6C, S6D, and S6F). Additionally, *ASTL,* which encodes ovastacin, a cortical granule protease that cleaves ZP2^30^, was expressed at a low level in GCs with a Hominini-enriched pattern (Figure S6C). Further validation by immunohistochemistry confirmed that ZP3 and ZPBP proteins are upregulated in humans and localized within cerebellar glomeruli alongside ZP2 protein (Figures 3E and 3F). ASTL protein was detected in GCs as well as astrocytes that wrap around the cerebellar glomeruli in all species, with similar localization to ZP2 (Figures S6G and S6H). Despite the well-characterized roles of ZP family members and ZPBP in reproduction, their potential function in the brain remains unexplored.

These findings prompted us to further examine the expression patterns of additional genes involved in oocyte-sperm interaction. Gene lists were compiled based on GO terms associated with the zona pellucida receptor complex, sperm-egg recognition, as well as sperm and acrosomal membrane. Remarkably, many of these genes were expressed in the cerebellum across species (Figure S6E). Notably, among them, genes related to sperm membrane (*ASAH1*, *MSR1*, and *ANKUB1*), acrosomal membrane or matrix (*CTNNA1*, *DSE*, *ANXA11*, and *SPESP1*), spermatogenesis and spermiogenesis (*CXADR*, *RACGAP1*, and *MAEL*), and response to estrogen (*CFLAR* and *ESR2*) exhibited human-specific upregulation in GCs (Figures 3C and S6F). The expression of ZP family members, their putative interactors, and gamete interaction-related genes in GCs, which are neurons within the cerebellum, hints at a potentially novel, human-specific function or regulatory mechanism distinct from their known role in reproduction.

### Gene Regulatory Network Predicting Regulation of Human-Specific ZP2 Expression

To explore the regulatory mechanisms that drive *ZP2* expression, we first examined chromatin accessibility near the *ZP2* gene locus (Figure 4A). Prominent accessible peaks were detected in human GCs but were absent in other cell types or primates, indicating GC- and human-specific transcriptional activation of *ZP2* (Figures 4A, S7A, S7B). Consistent peaks across four human donors (Figure S7C) further support the robustness and reproducibility of these signals. Interestingly, adjacent genes *CRYM, TMEM159,* and *ANKS4B* also exhibited human-enriched chromatin accessibility and upregulation in GCs (Figure 4A, S7B, Table S1). This genomic region, spanning *CRYM*, *ZP2*, and *TMEM159,* has been previously reported to show human-specific co-regulation^25,36^. Consistently, our single-cell transcriptomic analysis revealed that these genes show similar expression patterns during development and co-expression with ZP2 in specific GC subpopulations (Figures S7D and S7E). Using SCENIC+, we predicted gene regulatory networks and identified eight putative regulatory regions that may regulate *ZP2* transcription. (Figure 4B). Among these *ZP2* regulatory regions, three were also predicted to regulate *CRYM* and *TMEM159*, further indicating their coordinated regulation through shared regulatory elements. Moreover, we found that two ZP2 promoter regions overlap with cerebellum-specific expression quantitative trait loci (eQTLs) for *ZP2* (https://www.gtexportal.org/), supporting our predicted regulatory elements and suggesting that genetic variation in these regions may contribute to ZP2 regulation in humans.

**Figure 4.**
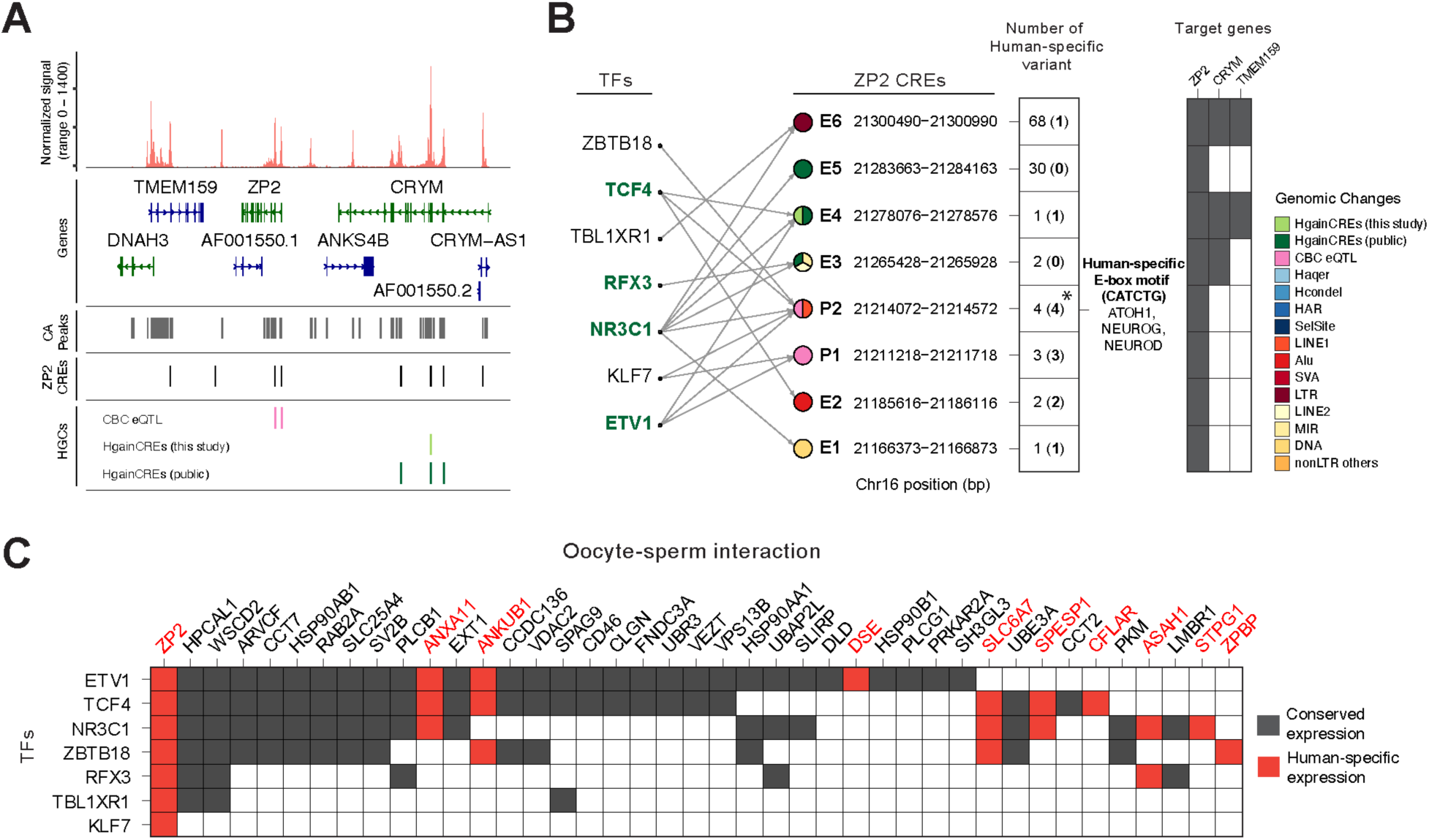
Gene regulatory network and cis-regulatory sequence differences predicted to drive human-specific *ZP2* expression. (A) Chromatin accessibility profiles surrounding the ZP2 gene loci in GCs of the adult human cerebellum. HgainCREs: human-gained cis-regulatory elements identified in this study (this study) and from public datasets (public)^38^. ZP2 CREs: peaks associated with *ZP2* identified through gene regulatory network analysis using SCENIC+. Cerebellum-specific expression quantitative trait loci (CBC eQTL) data were obtained from the GTEx Portal (https://www.gtexportal.org/). (B) Predicted transcription factors (TFs) and putative regulatory elements associated with ZP2 expression. Various genomic modifications were examined, including human-gained cis-regulatory elements (HgainCREs)^38^, human ancestor quickly evolved regions (Haqer)^82^, human-specific conserved deletions (Hcondel)^83^, human accelerated regions (HAR)^84–89^, and natural selection sites (SelSite)^90–92^. Repeat elements were obtained from the UCSC genome browser (https://genome.ucsc.edu/cgi-bin/hgTables) and analyzed, including long interspersed nuclear elements (LINE), Alu elements (Alu), long terminal repeats (LTR), SINE-VNTR-Alu retrotransposons (SVA), microRNA (MIR), and other transposable elements (nonLTR others). TFs predicted to bind to the *ZP2* CREs that contain HgainCREs are highlighted in bold. The number of human-specific sequence variants identified in each CRE relative to Hominidae is shown, with unfiltered counts and filtered counts (in bold). Filtered counts exclude nearby, low-confidence, or INDEL-associated variants. *, A human-specific sequence variant was identified in the promoter region P2, generating a high-affinity E-box motif recognized by proneural bHLH TFs. (C) Integrative gene regulatory network predicting TFs regulating *ZP2* and oocyte-sperm interaction genes. Genes showing human-specific expression are highlighted in red. See also Figure S7.

These ZP2 regulatory regions are predicted to be bound by seven TFs: *ZBTB18*, *TCF4*, *TBL1XR1, RFX3, NR3C1, KLF7,* and *ETV1* (Figure 4B). These TFs exhibit developmental expression patterns closely aligned with *ZP2* expression, supporting their potential role in *ZP2* regulation (Figure S7F). To further define the regulatory elements responsible for human-specific *ZP2* expression, we identified human-gained cis-regulatory elements (HgainCREs) based on chromatin accessibility patterns in our single-cell data from four primates. Additionally, we analyzed overlaps with previously reported human- and cerebellum-specific CREs based on ChIP-seq datasets from humans, chimpanzees, and macaques^38^. Three of the *ZP2* regulatory regions contain HgainCREs, suggesting their potential role in human-specific *ZP2* activation. Furthermore, we identified that *ETV1*, *NR3C1*, *RFX3*, and *TCF4* bind to these HgainCREs-associated regions (Figure 4B) and exhibit higher expression levels in the cerebellum compared to other brain regions (Figure S7G). Notably, *ETV1* has been shown to regulate activity-dependent synaptic maturation in cerebellar GCs^39^, while *NR3C1*, encoding a glucocorticoid receptor, modulates synapse plasticity and neuronal remodeling through its role in stress-related circuit regulation^40^.

We next asked whether these regulatory elements harbor sequence changes unique to humans. Comparative genomic analysis can identify human-specific sequence variants that alter TF binding or cis-regulatory activity, providing a potential mechanism for the human-specific activation of ZP2. We performed comparative sequence analysis of the eight candidate CREs identified (Figure 4B) using multiple sequence alignments spanning 447 mammalian species^41–44^. We quantified sequence differences between humans and distinct phylogenetic groups, including Hominidae (chimpanzees, bonobos, gorillas, orangutans), Hylobatidae, Cercopithecoidea, Platyrrhini, Tarsiiformes, Strepsirrhini, and all mammals (Figures S7H and S7I, Table S2). The distribution of human-specific variants varied across CREs. Although CREs E5 and E6 showed the highest density of changes, these regions are enriched for repetitive and long terminal repeat (LTR) elements, likely reflecting their divergence. After excluding low-confidence substitutions (see Methods), we identified 12 sequence variants unique to humans relative to other Hominidae, of which 4 were unique relative to all primates and 1 remained unique across all mammals (Figures 4B and S7I).

To assess whether these human-specific variants could affect TF binding, we intersected the 12 variants with TF binding site predictions using MotifbreakR. Only one variant created a strong binding site in the promoter region P2, corresponding to an ultraconserved position across mammals. This variant generates an E-box motif (CATCTG) preferred by proneural bHLH factors, including NEUROG, NEUROD, and ATOH1^45^. ATOH1 is a key regulator that specifies all rhombic lip-derived GC lineages during cerebellar development^46^. While ATOH1 expression declines, ZP2 expression increases postnatally, with overlap in early postnatal stages. This raises the possibility that the human-specific variant in P2 enables early ATOH1 binding to ZP2 regulatory elements in GCs. As development progresses, other CREs may be engaged by TFs identified in our GRN analysis (Figure 4B), together contributing to the human- and cerebellum-specific expression of ZP2. These findings suggest that human-specific sequence and chromatin accessibility changes within ZP2 regulatory regions may modulate TF binding, contributing to its human-specific and cerebellum-specific expression. Collectively, these human-specific CREs and predicted TF interactions represent promising candidates for future experimental validation to uncover upstream drivers of ZP2 regulation.

To investigate the potential involvement of convergent upstream mechanisms in the expression of *ZP2* and other oocyte-sperm interaction genes in the human cerebellum, we examined whether the seven TFs predicted to bind *ZP2* regulatory regions also target these genes (Figure 4C). Our analysis revealed that several targets, including nine with human-specific expression, are predicted to be regulated by the same TFs, suggesting a coordinated transcriptional network. This shared regulation may represent an evolutionary repurposing of gene regulatory programs in the human cerebellum, extending beyond their described roles in reproduction.

### ZP2 Protein Localizes at Cerebellar Synaptic Glomeruli

Given the robust evidence of human-specific expression of ZP2 and its putative interactors in GCs, we investigated its potential cellular function in the cerebellum. To explore whether *ZP2* expression correlates with specific cellular states or molecular signatures, we assessed the differentially expressed genes (DEGs) between *ZP2*-positive and *ZP2*-negative populations in our adult cerebellum dataset. This analysis revealed few DEGs, possibly due to the low *ZP2* expression levels or its role as a secreted molecule. We then subclustered the developmental datasets^17^ by specific periods and found significant DEGs during postnatal periods (Figure 5A). Upregulated genes in *ZP2*-positive GCs were enriched for GO terms associated with synapse activity regulation, axon guidance, and Rho GTPase and RHOBTB3 signaling. Downregulated genes were associated with cell junction organization, ROBO signaling, and synapse organization regulation. Since *ZP2* encodes a secreted protein, these transcriptional differences likely reflect a shared regulatory network controlling both *ZP2* and synaptic genes, highlighting their strong transcriptional correlation.

**Figure 5.**
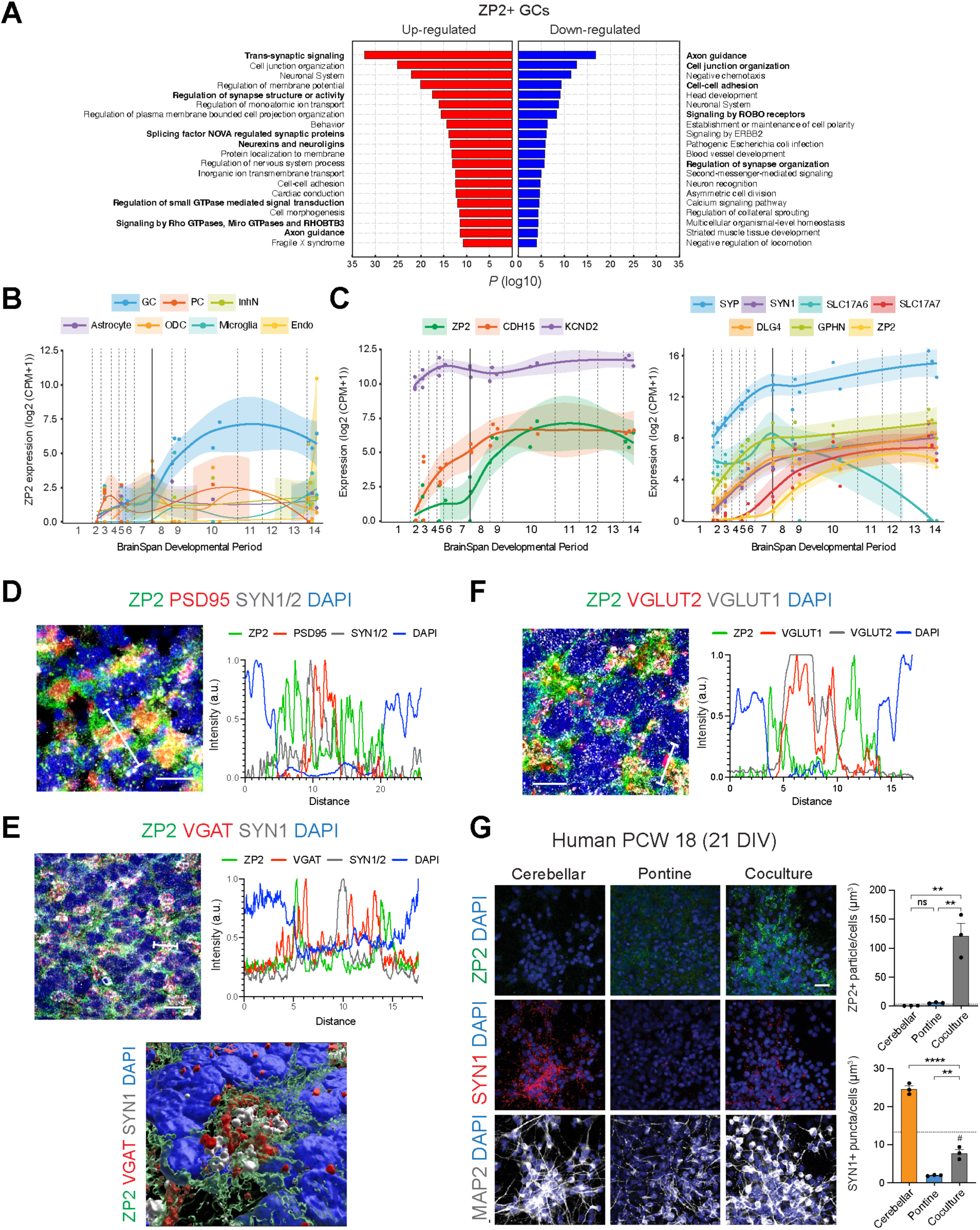
Human-specific expression of ZP2 at pontocerebellar glomerular synapses. (A) Enriched Gene Ontology (GO) terms for differentially expressed genes between *ZP2*-positive and *ZP2*-negative cells in the human postnatal cerebellum^17^. (B) *ZP2* expression across cell types during development, depicted by pseudo-bulk RNA-seq analysis of developing human cerebellum data^17^. Values represent individual donors. (C) Developmental expression patterns of glomerulus maturation regulators *CDH15* and *KCND2*, excitatory synaptic protein genes *SYN1*, *SYP*, and *DLG4/PSD95*, inhibitory synaptic protein genes *GPHN*, and glomerular presynaptic markers *SLC17A7/VGLUT1* and *SLC17A6/VGLUT2* in GCs^17^. Values represent pseudo-bulk expression levels from individual donors. (D-F) Spatial distribution of ZP2 and excitatory synaptic proteins SYN1 and PSD95 (D), inhibitory synaptic protein VGAT (E), glomerular presynaptic markers VGLUT1 and VGLUT2 (F) within cerebellar glomeruli. The spatial plot shows representative protein signal intensities along the marked line in the images. A 3D reconstruction of the ZP2-VGAT-SYN1 signals further illustrates their close spatial relationship within glomerular structures. (G) Immunohistochemistry of human cerebellar, pontine, and cerebello-pontine co-culture obtained at post-conception week (PCW18) and cultured for 21 days in vitro, with quantification of ZP2 particle and SYN1 synaptic puncta volume. n=3, technical replicates. Statistical comparisons performed with Student’s t-test, p-values: ns – not significant, ** < 0.01 and **** < 0.0001. See also Figure S8.

Reminiscent of the zona pellucida, the cerebellar glomerulus is surrounded by an extracellular matrix and a glial sheath that anchor synapses^4,47^. Based on the specific expression of ZP2 in GCs and the highly regulated connectivity between GC dendrites and mossy fiber axons, we hypothesized that ZP2 may play a role analogous to its function in the zona pellucida, potentially regulating synapse organization among GCs, inhibitory interneurons Golgi cells, and incoming mossy fibers. The developmental expression pattern of *ZP2* (Figure 5B) also supports its possible involvement in synaptic functions. *ZP2* expression in GCs begins around birth, increases progressively postnatally, and declines slightly with aging^48^. This expression pattern is closely followed by *CDH15* and *KCND2,* genes associated with cerebellar glomerular maturation^49,50^, and peaks with other synaptic genes (Figure 5C). Immunohistochemical analysis of the adult human cerebellum demonstrated distinct localization of ZP2, CDH15, and KCND2 within glomeruli (Figures S8A and S8B). Notably, ZP2 was predominantly localized at the periphery of CDH15/KCND2 signals. Additionally, ZP2 signals were negatively correlated with excitatory and inhibitory synaptic proteins, as well as with other glomerular presynaptic markers VGLUT1 and VGLUT2, which are associated with mossy fiber terminals from dorsal column nuclei and spinocerebellar tracts, respectively^51^ (Figures 5D-F and S8B-D). Spatial analysis revealed that excitatory synaptic proteins were concentrated centrally, surrounded by inhibitory synaptic proteins, while ZP2 signals were highest at the periphery of these synaptic proteins within the glomerulus structure (Figure 5E). This distinctive localization suggests ZP2’s potential influence on synaptic composition and functionality within cerebellar glomeruli.

Considering the correlation between the developmental expression pattern of *ZP2* and synapse maturation in cerebellar glomeruli, we hypothesized that incoming pontine mossy fibers, known to initiate glomerulus formation and maturation^52–55^, might trigger ZP2 expression in GCs. We tested this using human cerebellar and pontine cells isolated at post-conception week (PCW) 18, a stage before the onset of ZP2 expression (Figure 5G). After validating the expression of canonical markers of the cerebellum (ZIC1) and pons (HOXA2, HOXB4)^53^ (Figure S8E), cerebellar cells were cultured either alone or with pontine cells. After 21 days, ZP2 protein levels were significantly elevated in co-cultures, whereas cerebellar or pontine cells cultured alone expressed little to no ZP2 (Figure 5G). Furthermore, co-cultures showed significantly lower levels of the presynaptic protein SYN1 compared to cerebellar cells cultured alone. This ZP2 induction in co-culture was consistent across cultures from different donors, with a concurrent reduction in another presynaptic protein SYP (Figure S8F). These findings suggest that the projection of pontine mossy fibers into the cerebellar cortex triggers ZP2 expression in GCs, which subsequently modulates and restricts synaptogenesis.

### ZP2 Regulates Synapse Development and Neuronal Activity

To investigate whether ZP2 directly regulates synapse formation, we treated cerebellar and pontine cells with recombinant ZP2 protein (Figures 6A and S9A). Treatment with human ZP2 significantly reduced SYN1-positive synaptic puncta in cerebellar cells, similar to the effect observed in cerebellar-pontine cell co-cultures. In line with previous findings^56^, co-culture increased PSD95-positive postsynaptic puncta (Figure 6A). However, the addition of exogenous ZP2 protein attenuated this increase, reducing synaptic puncta colocalized with SYN1 and PSD95 (Figure S9A). These results support a model in which pontine input promotes postsynaptic compartment development while simultaneously inducing ZP2 expression as a feedback mechanism to limit excessive synapse formation.

**Figure 6.**
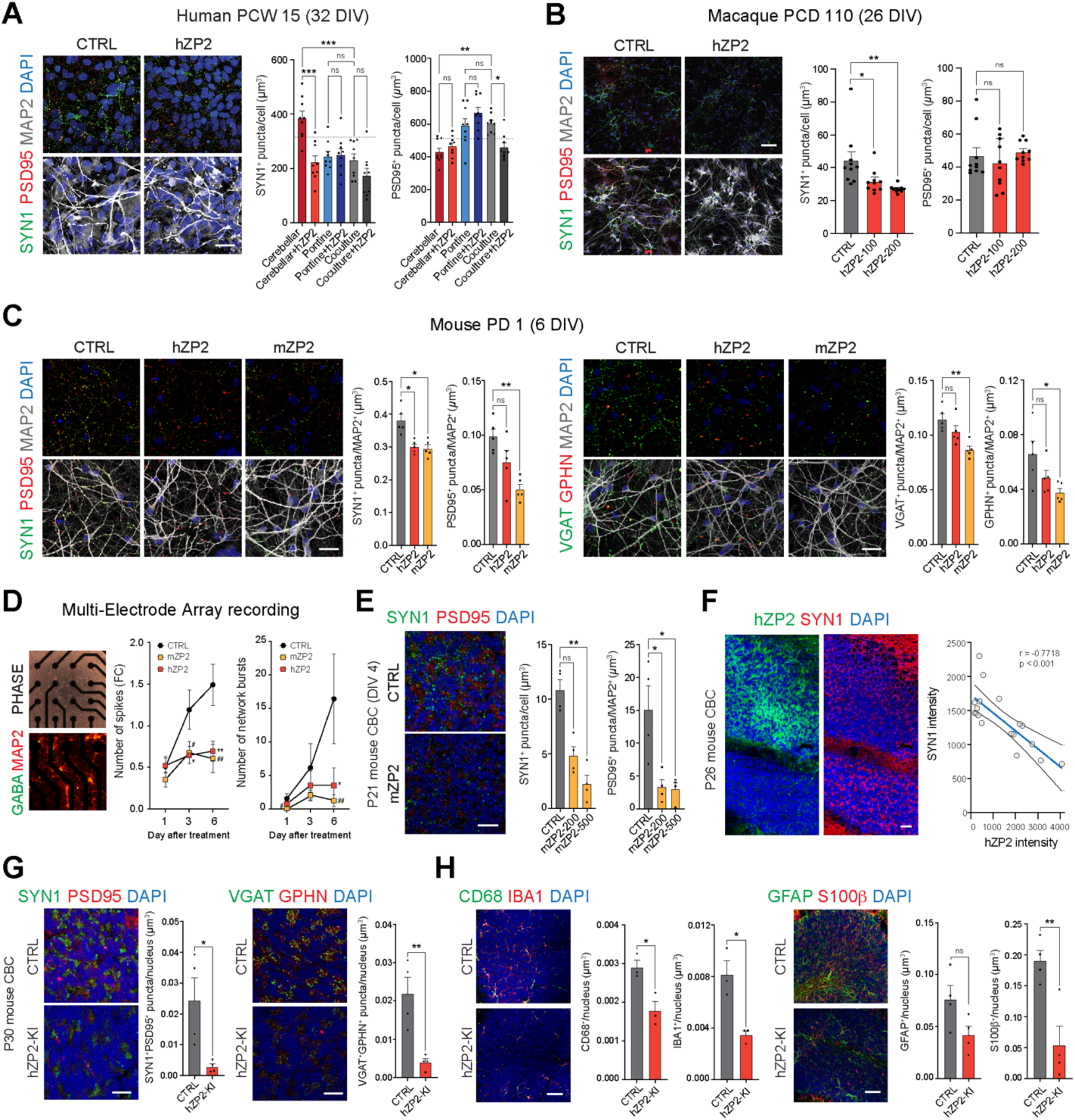
Human-specific regulation of synaptic development and neuronal activity by ZP2. (A) Reduced synaptic protein expression in human cerebellar cells after 48 hours of treatment with human ZP2 protein. Representative images and quantification of synaptic puncta volume after treatment. n=9, multiple fields from 3 technical replicates. (B) Reduced synaptic protein expression in macaque cerebellar cells (embryonic day E110) after 48 hours of human ZP2 treatment. n=10, multiple fields from 2 technical replicates. (C) Reduced excitatory (SYN1 and PSD95) and inhibitory (VGAT and GPHN) synaptic puncta in mouse cerebellar cells (postnatal day 1) after 48 hours of treatment with mouse and human ZP2. n=5, independent experiments. (D) Multi-electrode array recordings showing total spikes and network bursts normalized to pre-treatment. n=15, technical replicates. (E) Reduced SYN1 and PSD95 synaptic puncta in postnatal day 21 mouse cerebellar tissue cultured for 4 days with 48 hours of mZP2 treatment. n=4, technical replicates. (F) Reduced SYN1 expression in postnatal day 26 mouse cerebellum, 5 days after intracranial injection of human ZP2. (G) Reduced excitatory (SYN1 and PSD95) and inhibitory (VGAT and GPHN) synaptic puncta at the cerebellar glomeruli in control (CTRL) and hZP2 knock-in (hZP2-KI) mouse cerebellum at postnatal day 30 (P30). n=4 mice per group. (H) Reduced microglial (CD68 and IBA) and astrocytic markers (GFAP and S100β) in CTRL and hZP2-KI mouse cerebellum at P30. n=4 mice per group. All statistical comparisons performed with Student’s t-test, p-values: ns – not significant, Control vs. hZP2; WT vs. hZP2+/+; * < 0.05, ** < 0.01, *** < 0.001, and **** < 0.0001. Control vs. mZP2; ^#^, < 0.05, ^##^ < 0.01, and ^###^ < 0.001. Scale bar, (A-E, G, H) 20 µm, (F) 40 µm. See also Figures S9 and S10.

To test if ZP2 regulation of synapse formation relies on human-enriched binding partners and/or functions analogously to its species-specific oocyte-sperm recognition^57^, we treated macaque and mouse cerebellar cells with human ZP2 protein. Remarkably, both species showed a reduction in SYN1-positive synaptic puncta, indicating the presence of necessary machinery for ZP2-mediated synaptic downregulation (Figures 6B, 6C, and S9B-D). This reduction in SYN1-positive presynaptic puncta was also dose-dependent (Figure 6B). Notably, the reduction in PSD95-positive postsynaptic puncta was not statistically significant in macaque and mouse cultures with human ZP2 (Figures 6B, 6C, S9B, and S9C). However, a more pronounced reduction in SYN1, SYP, and PSD95 puncta was observed in mouse cells treated with mouse ZP2 protein compared to those treated with human ZP2 protein (Figures 6C and S9D), suggesting some degree of species-specificity in this ZP2 function. Importantly, there was no significant change in total cell numbers or MAP2-positive area, indicating that the reduced synaptic puncta were not due to neuronal cell death or reduced neuronal processes (Figure S9E). Additionally, inhibitory synaptic proteins VGAT and GPHN were reduced by mouse ZP2 treatment, suggesting that synapses between GCs and inhibitory interneurons were also compromised (Figures 6C and S9C). These effects were consistently observed with prolonged ZP2 treatment, which led to enhanced reductions, and were not associated with changes in neuron numbers (Figure S9F).

To assess the functional consequences of the observed synaptic protein reduction, we performed multi-electrode array recordings on mouse cerebellar cells (Figures 6D and S10). Both mouse and human ZP2-treated cells showed significantly reduced electrical firing, with more pronounced effects from mouse ZP2 protein. To further investigate ZP2’s role in more mature cerebellar cells, we examined its impact on the developing mouse cerebellum during a stage when GC migration from the external to the internal germinal layer was mostly complete and synaptic maturation was actively occurring^2,8^. In organotypic cultures of postnatal day 21 cerebellar tissue, treatment with mouse ZP2 protein caused a dose-dependent reduction in SYN1- and PSD95-positive synaptic puncta (Figures 6E and S9G). Furthermore, intracranial injection of human ZP2 protein into postnatal day 26 cerebellum reduced SYN1-positive puncta, with SYN1 intensity showing a clear negative correlation with the intensities of injected human ZP2 protein (Figure 6F). These findings indicate that ZP2 is co-opted to regulate cerebellar synapse development and neuronal activity.

To address the absence of intact pontocerebellar circuitry in in vitro assays and better evaluate ZP2 function in a more physiological setting, we generated a human ZP2 knock-in (hZP2 KI) mouse model in which ZP2 expression is driven by the *Atoh1* promoter, enabling GC-specific expression from embryonic stages and allowing assessment of its long-term effects during cerebellar development. Analysis at postnatal day 30, a period of active circuit maturation, revealed a significant reduction in both excitatory and inhibitory synaptic puncta in the GC layer of hZP2 KI mice compared to controls (Figure 6G). To assess whether these synaptic changes were accompanied by altered glial involvement, we examined astrocytic and microglial markers in the cerebellar GC layer. In hZP2 KI mice, we found a significant reduction in IBA1, CD68, and S100β expression, with a modest, non-significant decrease in GFAP (Figure 6H). Microglial and astrocytic engagement is driven by robust synaptic cues and activity during postnatal development, and reductions in neuronal activity or synapse formation can suppress glial recruitment and functional responses^58,59^. In line with these, our in vitro data demonstrate that ZP2 reduces both spontaneous firing and synapse formation, supporting the interpretation that attenuated glial activation in hZP2 KI mice reflects reduced synaptic engagement. Together, our in vitro, ex vivo, and in vivo findings consistently demonstrate ZP2’s role in limiting synapse formation. Notably, the hZP2 KI mouse model provides direct and physiologically relevant evidence that GC-specific ZP2 accumulation at the cerebellar glomerulus restricts synapse development and glial engagement during postnatal circuit refinement.

### Cerebellar Expression of Disease Risk Genes

Cerebellar dysfunction is associated with motor deficits, such as impaired balance and coordination, commonly observed in neurodegenerative disorders like ataxia. Some, such as spinocerebellar ataxia, result from single-gene mutations. Beyond motor control, the cerebellum also contributes to cognitive and emotional processes and is increasingly linked to neuropsychiatric disorders^60^. Using our cerebellar dataset alongside equivalent prefrontal cortex (PFC) data^24^, we performed a systematic, cell-level comparison of gene enrichment patterns associated with both neurodegenerative and neuropsychiatric disorders across these regions (Figures 7A and 7B). We applied a gene fraction enrichment score (GFES) to assess relative enrichment across gene lists of varying lengths. GFES estimates enrichment based on the proportion of disorder-associated gene reads per nucleus and applies ANCOVA for comparisons, with cell type cluster identity as an independent categorical variable. We then compared ANCOVA regression estimates across cell types, disorders, and species.

**Figure 7.**
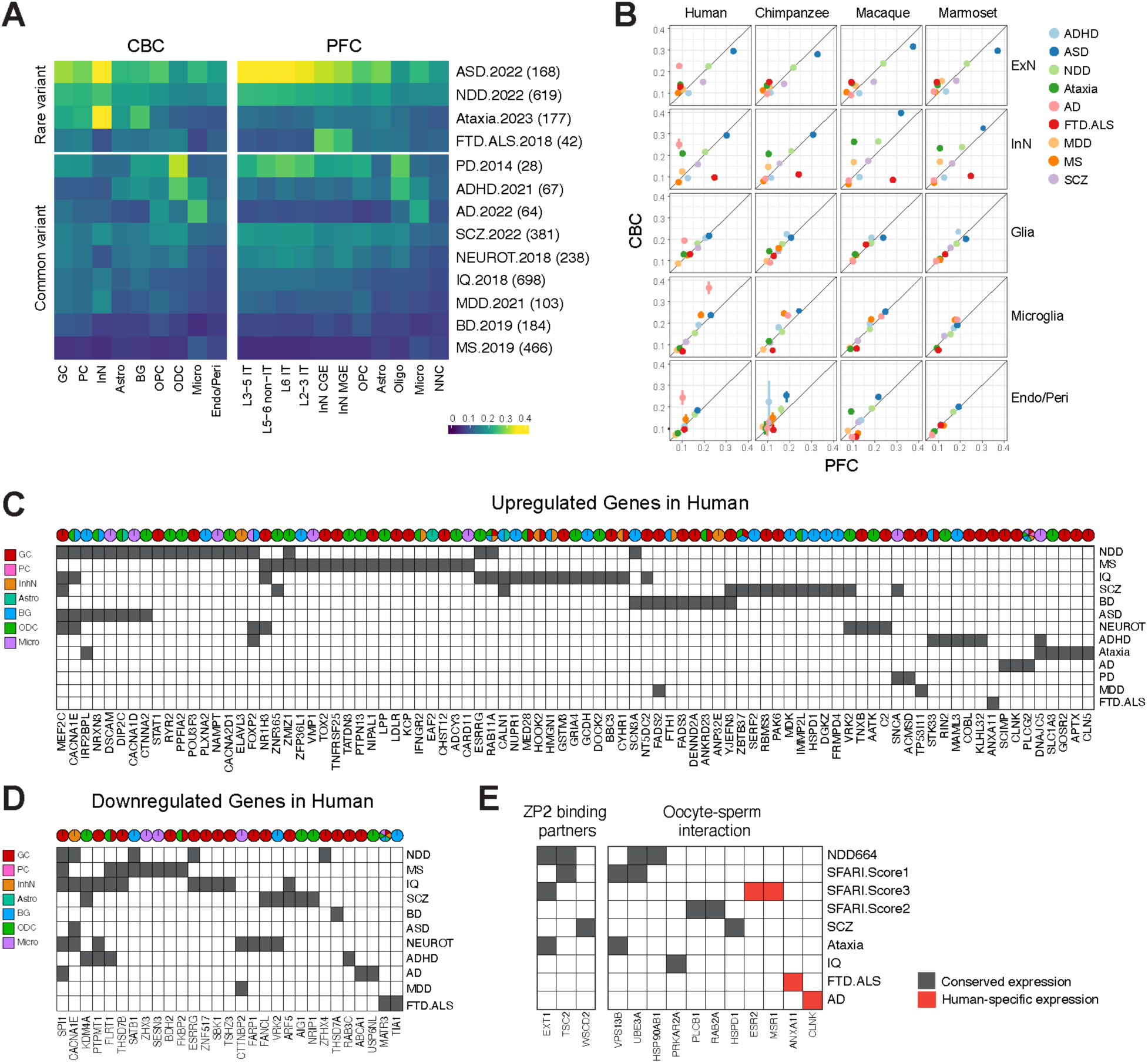
Human cerebellar association with brain disorders. (A) Enrichment of genes implicated in various neuropsychiatric and neurodegenerative disorders across cell types in the adult cerebellum (CBC) and prefrontal cortex (PFC). The numbers of genes for each disease category are indicated. (B) Correlation of risk gene enrichment between adult cerebellum and PFC across cell types and primate species. (C-D) Genes implicated in brain disorders exhibiting human-specific upregulation (C) and downregulation (D) across cerebellar cell types. (E) Genes encoding putative ZP2 binding partners and those involved in oocyte-sperm interaction implicated in brain disorders, including autism spectrum disorder (ASD, SFARI Score 1, 2, 3), neurodevelopmental disorders (NDD), ataxia, frontotemporal dementia and amyotrophic lateral sclerosis (FTD.ALS), Parkinson’s disease (PD), attention-deficit/hyperactivity disorder (ADHD), Alzheimer’s disease (AD), multiple sclerosis (MS), schizophrenia (SCZ), neuroticism (NEUROT), intelligence quotient (IQ), major depressive disorder (MDD), bipolar disorders (BD). See also Figure S11.

To validate our approach, we assessed risk genes with previously defined regional associations: 177 genes linked to ataxia and 42 genes associated with frontotemporal dementia (FTD) and amyotrophic lateral sclerosis (ALS). As expected, ataxia-related genes were highly enriched in cerebellar InNs. Notably, FTD-ALS genes showed high enrichment in PFC InNs, a cell-type specificity that has been less extensively characterized (Figures 7A and 7B). For ASD-associated genes, we observed substantial enrichment in multiple subtypes of excitatory neurons in the PFC, with less enrichment in the cerebellum, consistent with the primarily cortical involvement of these cell types in ASD. Inhibitory neurons, however, displayed similar enrichment for ASD-associated genes in both the cerebellum and cortex. Genes linked to neurodevelopmental delay (NDD) showed modest enrichment. This aligns with previous studies suggesting that NDD-associated genes act earlier in development compared to ASD-associated genes^61^, which may explain the lower enrichment in our analysis of adult tissues.

We next examined gene sets derived from common variant studies, specifically from genome-wide association studies (GWAS). GWAS identifies genomic loci associated with traits or diseases; however, linking these loci to specific genes remains challenging. Consequently, GWAS-identified genes are often more challenging to link directly to disease mechanisms compared to rare variants, resulting in less pronounced gene enrichment estimates. The enrichment pattern for SCZ-associated genes resembled that of ASD, with strong enrichment in cortical excitatory and inhibitory neurons in both the PFC and cerebellum. Alzheimer’s disease (AD)-associated genes showed the highest enrichment in microglia (Figures 7A and 7B). Interestingly, we observed substantially greater microglial enrichment in the cerebellum, which appeared to be human-specific. Further analysis revealed that this enrichment was primarily driven by a small nuclear pseudogene *RN7SKP176*, which showed human-specific upregulation and overlaps with the 3‘UTR region of the AD-associated gene *PLCG2* (Figure S11). While it remains unclear whether *RN7SKP176* has a role in AD pathogenesis, this pseudogene may interfere with eQTL co-expression analysis, warranting further investigation.

We further identified disease-associated risk genes (https://omim.org, https://gene.sfari.org, see Methods) with human-specific expression patterns in the cerebellum at the cell type level to examine how evolutionary transcriptomic changes may intersect with genes relevant to brain disorders. (Figures 7C and D). Among these, *CACNA1E*, implicated in ASD and NDD, exhibited human-specific upregulation in BGs and ODCs, along with downregulation in InNs. Similarly, *MEF2C*, also associated with these disorders, showed human-specific upregulation in GCs. Building on our findings of ZP2’s role in synapse formation and neuronal activity, we further examined whether genes encoding putative ZP2 interactors and those involved in oocyte-sperm interaction are also implicated in brain disorders (Figure 7E). Several of these genes have been previously associated with brain disorders. Notably, ZP2 binding partners *EXT1*, *TSC2*, and *WSCD2* have been associated with NDD, ASD, SCZ, and ataxia. *ESR2* and *MSR1*, which exhibit human-specific expression, have been implicated in ASD, while *ANXA11* has been linked to FTD and ALS. These analyses highlight brain disorder risk genes with human-specific expression and functions related to gamete interaction at the cell type level, offering potential new insights into the molecular mechanisms underlying these disorders and their function in the cerebellum.

## DISCUSSION

This study presents a comprehensive cellular dataset detailing gene expression and regulatory networks in the adult primate cerebellar cortex, providing a unique resource for investigating conserved, divergent, or uniquely human molecular features across primates. Leveraging this dataset, we uncovered novel biological insights into human-specific cerebellar characteristics. We found that human GCs exhibit the highest transcriptomic divergence, reflecting significant evolutionary specialization. GO analysis revealed that human and Hominini-specific expression is prominently associated with synaptogenesis, axon elongation, and synapse maturation. Notably, several genes associated with oocyte-sperm recognition, the sperm membrane, and spermatogenesis are specifically upregulated in human GCs, suggesting a potential evolutionary repurposing of reproductive genes in the human cerebellum. Our findings illustrate the intricate interplay of genetic, evolutionary, and developmental processes that shape the human brain.

Among the genes exhibiting human-specific expression patterns, *ZP2* was the most specific to the human cerebellum. Its developmental expression patterns, as well as our *in vitro* findings using cerebellar and pontine cell co-cultures, suggest that ZP2 expression in GCs may be regulated by incoming pontine mossy fibers during development. Additionally, our findings demonstrate that ZP2 protein reduces excitatory and inhibitory synaptic puncta, likely inhibiting synapse formation among GCs, mossy fibers, and Golgi cells at the glomeruli. This is further supported by its peak expression following glomerular maturation and its distinct localization within cerebellar glomeruli. ECM components play critical roles in regulating synapse formation, plasticity, and neuronal activity^62^. Extending these observations, we describe a novel function of the ECM protein ZP2 in restricting excitatory and inhibitory synapse formation at the glomeruli, potentially regulating human pontocerebellar circuitry. Notably, major components of ZP filaments, including ZP1-3, are expressed in human GCs, and the protease ovastacin, which cleaves ZP2 and initiates its polymerization, is expressed in both GCs and astrocytes. This raises the possibility that ZP2 may regulate synapse formation through a mechanism similar to the recently described ZP filament cross-linking process, which rigidifies the egg coat^31^. Importantly, ZP3 expression is undetectable or extremely limited in the mouse cerebellum^15,17^, yet both recombinant ZP2 treatment of mouse cultures and human ZP2 KI mice revealed significant synapse reduction. These findings suggest that ZP2 can act independently of ZP3 to regulate synaptic development. Further investigation is needed to determine whether ZP2 polymerizes with other ZP family proteins in the human cerebellum and how this process influences ECM properties and synapse formation.

While genes typically associated with oocyte-sperm recognition are expressed in the human cerebellum and, in the case of ZP2, influence synapse development and neuronal activity, the extent to which these phenotypic changes confer adaptive advantages for human fitness remains uncertain. The cerebellum of humans and great apes (hominids) has undergone significant expansion, particularly in the posterolateral regions, which receive projections from the reciprocally expanded prefrontal and posterior associative cortices^63,64^. As the cerebellum enlarges, GC numbers increase disproportionately relative to pontine mossy fibers, leading to greater synaptic load and structural complexity at cerebellar glomeruli^4^. This expanded GC pool relative to pontine inputs may place increased demands on synaptic refinement, potentially necessitating additional regulatory mechanisms such as ZP2-mediated synaptic modulation. Although the mechanisms driving these evolutionary adaptations in connectivity remain elusive, we propose that human-specific ZP2 expression slows the development of pontocerebellar glomerular synapses and suppresses premature synapse formation, thereby extending the period for circuit assembly and reorganization. This prolonged developmental window, reflecting a retention of neotenic features, may allow for the coordinated formation of increasingly complex glomeruli in the human cerebellum, ultimately supporting fine motor and cognitive functions. Notably, evidence indicates that humans acquire fine motor skills later than other primates, potentially due to their larger brains requiring longer periods for developing dexterity, which partially relies on cerebellar circuits^65^. Previous studies have revealed evolutionary changes in human synapse development and identified human-specific synaptic modifiers in the cerebral cortex^16,66–68^, but similar regulators have not been described in the cerebellum.

The human cerebellum is densely populated with GCs, which undergo prolonged neurogenesis during early postnatal development^18,69,70^. Approximately 85% of the GCs are generated postnatally in humans, with the GC-to-PC ratio increasing from 485:1 in the first postnatal month to 2,700:1 by 10-11 months^70^, eventually reaching an adult ratio of roughly 3,000∼5000:1^71,72^. On the other hand, neurons in the basal pons are generated and begin extending axonal projections to the cerebellum much earlier^53^. This extended generation of GCs poses significant challenges in establishing precise connectivity with incoming pontine mossy fibers. Secreted molecules such as WNT7A and FGF22 promote pontocerebellar synaptogenesis^73,74^. Additionally, GCs in the external germinal layer and CDH7 expression in GCs of the internal germinal layer restrict mossy fiber extension into the proliferative regions during development^54,55^. In addition to these regulatory mechanisms, our findings suggest that ZP2 serves a crucial function in limiting synapse formation in pontocerebellar connections, highlighting its significance as an evolutionary adaptation associated with the expanded GC population and prolonged maturation of the human cerebellum.

Moreover, the decreased neural activity in GCs upon exposure to ZP2 protein suggests that ZP2 may also participate in adaptive regulation to reduce the energy demands of the expanded human GC pool. If each GC consumed energy at levels comparable to typical cortical neurons, the metabolic demands would likely be unsustainable. Notably, GCs generally exhibit relatively low firing rates compared to PCs and cortical pyramidal neurons^4,5^. While our findings support a role for ZP2 in limiting synapse formation during development, it remains to be determined whether ZP2 modulates the timing or complexity of glomerular maturation.

Given ZP2’s role in cerebellar glomerular synaptogenesis and neural activity, recent advances in human cerebellar organoid systems, which generate cerebellar-like structures with functional neurons^75,76^, provide a promising model to further explore ZP2’s role in development. However, current models remain limited in their ability to capture late postnatal stages and lack key afferent inputs such as pontocerebellar projections. With further refinement, these systems could serve as a powerful tool to dissect the mechanisms underlying ZP2 function in human cerebellar development and connectivity.

The ZP family and many gamete interaction genes are predominantly associated with infertility, and the species-specific sperm–oocyte binding depends on ZP2^57^. Rare homozygous mutations in ZP2, leading to truncated proteins that cannot be transported efficiently to zona pellucida (ZP), have been reported in women with a thin ZP and defective sperm binding to the oocyte^77–79^. However, no neurological assessment has been conducted in those studies to determine if those variants might also impact brain development and function. Intriguingly, KEGG analysis revealed that downregulated genes in rat oocytes engineered to carry the human infertility-causing variant of ZP2 were enriched in several signaling pathways, including those associated with glutamatergic synapses^79^. This suggests a potential shared dysregulation of genes between ZP2-dependent oocyte function and synapse activity. Although direct evidence connecting genetic variations in ZP family members and their interactors to human brain disorders is currently lacking, downregulation of *ZP2* has been observed in the cerebellum of patients with medulloblastoma, Huntington’s disease, or ASD^36,80^. Interestingly, we also observed a decline in ZP2 expression with aging in humans in this study, possibly related to the age-dependent reduction in glomerular synaptic contacts^81^. Although this study has certain limitations, future research with a broader and deeper examination of ZP2 and other gamete interaction genes will enhance our understanding of their potential contributions to human cerebellar development, evolution, function, and dysfunction.

### Limitations of the study

While this study provides valuable insights into human-specific gene regulation in the cerebellum, some limitations should be noted. The modest sample size across species, due to challenges in obtaining high-quality postmortem primate brain tissue, may affect the robustness of certain findings. Variability in tissue quality, genetic background, or environmental factors could also influence gene expression profiles. Transcriptomic resolution for rare cell populations, such as Purkinje cells, is limited in our dataset, reflecting both their biological rarity and high susceptibility to dropout in snRNA-seq. ZP2 function in synapse regulation and neural activity was examined with fetal human cerebellar and pontine cells in vitro and more robustly in mouse models. Further investigations of spatiotemporal dynamics of ZP2 expression in developing human pontocerebellar connections are needed to further validate and strengthen these findings. While ZP2 impacts both synapse integrity and neuronal activity, their causal relationship remains unclear and will require time-resolved, single-cell analyses capable of simultaneously tracking neuronal activity and synaptic changes.

## RESOURCE AVAILABILITY

### Lead contact

Further information and requests for resources and reagents should be directed to and will be fulfilled by the lead contact, Nenad Sestan.

### Materials availability

Unique reagents generated in this study are available from the lead contact upon completion of a Materials Transfer Agreement.

### Data and code availability

Processed data with relevant public datasets can be interactively explored at https://nemoanalytics.org/p?l=SestanPrimateCerebellum. All deposited data are publicly available as of the date of publication. Other relevant data and materials, as well as any additional information required to reanalyze the data reported in this paper, are available from the lead contact upon request.

## ACKNOWLEDGMENTS

This study was supported by NIH grant (MH123074-01) to A.C.; National Chimpanzee Brain Resource grant (NS092988), National Science Foundation grants (EF-2021785, DRL-2219759), and NIH grants (AG067419, HG011641) to C.C.S.; Center for Restoration of Nervous System Function grant (RX002999-01) to M.E. and S.G.W.; NIH grants (HD106197, MH130991), SFARI pilot grant, Brain and Behavior Research Foundation grant (29721), and Brain Research Foundation grant (BRFSG-2023-11) to A.M.M.S.; NIDA Merit Award DA023999 to P.R.; Medical Scientist Training Program grant T32 (GM140935) and Wisconsin Distinguished Graduate Fellowship to R.D.R.; AEI/10.13039/501100011033 grant (PID2019-104700GA-I00, PID2022-140137NB-I00), Fundació La Marató de TV3 grant (202230-30), and Instituto de Salud Carlos III (CP20/00064), with co-financing by European Funds for the Miguel Servet Contract to G.S.; NIH grants (HG012108, HG010898, HG012483, MH130991, U01MH116488, U01DA053628) to N.S.

## AUTHOR CONTRIBUTIONS

Conceptualization, S.K., A.C., and N.S.; Tissue procurement, S.P., A.H., M.G., Y.M.M., E.W.D., K.T.G., J.J.E., P.R.H., M.M.D., C.C.S., A.D., and A.M.M.S.; Multiome processing and library preparation, S.K., A.C., R.K., and S.P.; Computational analysis and data interpretation, S.K., A.C., S.M., Y.L. X.M, J.M. R-J, N.M., G.S., M.L., M.Z., D.M.Y., V.L., A.N., D.L., H.C., S.P., A.S., X.L., A.K., and C.C.; Immunohistochemistry and in situ hybridization, S.K., A.S., S.H., H.K., A.S., and A.M.M.S.; Tissue culture, S.K., A.C., S.H., H.K., and N.M.; Mouse intracranial injection, S.K.S., and I.C.; hZP2-KI mouse analysis, Y.Z., S.K., I.C.; Multi-electrode array recordings, N.M. and M.E.; Supervision, S.G.W., P.P., S.J.S., and N.S.; Project administration, N.S.; Writing for the original draft, S.K., A.C., and N.S.; Editing, S.K., A.C., and N.S., and all authors.

## SUPPLEMENTAL INFORMATION

**Document S1. Figures S1–S11**

**Table S1. Summary of cerebellar tissue used for Multiome analysis and gene lists with conserved and divergent expression patterns.**

**Table S2. GO and SynGO enriched terms for human-specific divergent genes and comparative sequence analysis of ZP2 CREs.**

## SUPPLEMENTAL FIGURES

**Figure S1.**
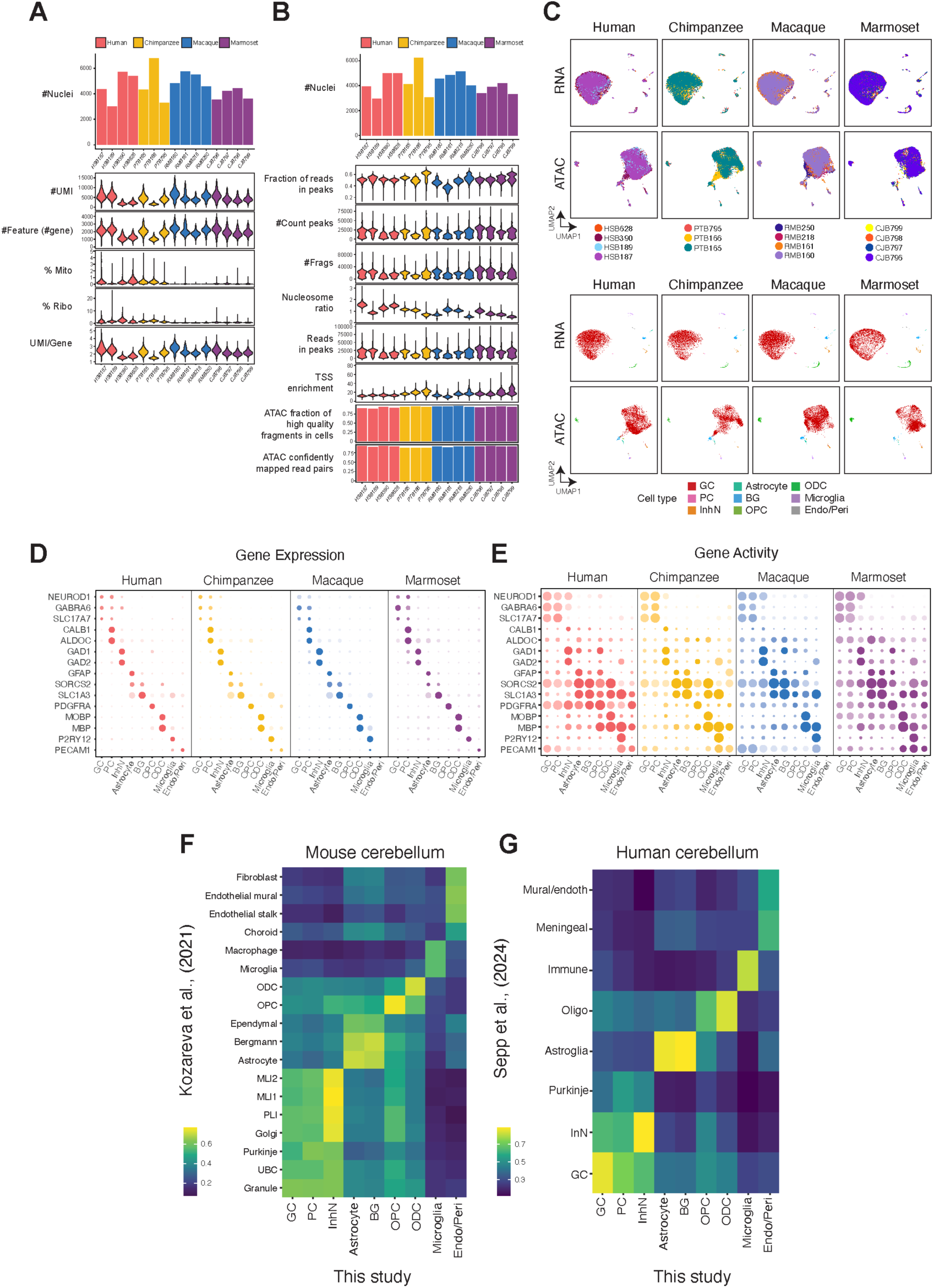
Quality control for single-nucleus Multiome data and cerebellar cell-type annotation, related to Figure 1. (A) A total of 69,302 high-quality nuclei were included for RNA-seq after quality control. Top: Bar plots indicate nuclei counts per sample. Bottom: Violin plots demonstrate the quality of the final dataset, showing the number of unique molecular identifiers (#UMI), number of genes (#Gene), percent mitochondrial reads (%Mito), and percent ribosomal reads (%Ribo) for each sample. (B) A total of 63,491 high-quality nuclei were included for ATAC-seq after quality control. Top: Bar plots indicate nuclei counts per sample. Bottom: Violin plots demonstrate the quality of the final dataset, showing the fraction of reads in peaks, number of peaks (#Peaks), number of fragments (#Frags), transcription start site (TSS) enrichment score, fraction of high-quality fragments in cells, and confidently mapped read pairs. (C) UMAP representations of cluster annotations for human, chimpanzee, macaque, and marmoset datasets, colored by donors and cell types. (D-E) Gene expression (D) and activity score based on chromatin accessibility (E) of canonical cell type-specific markers. (F-G) Spearman correlation of pseudo-bulk transcriptomes for cell type clusters between our dataset and adult mouse^15^ (F) or developmental human cerebellar scRNA-seq dataset^17^ (G). All major cell types except Purkinje cells show strong transcriptomic similarity across datasets. The weaker or ambiguous correlation of Purkinje cells likely reflects their low abundance and increased susceptibility to dropout in snRNA-seq datasets.

**Figure S2.**
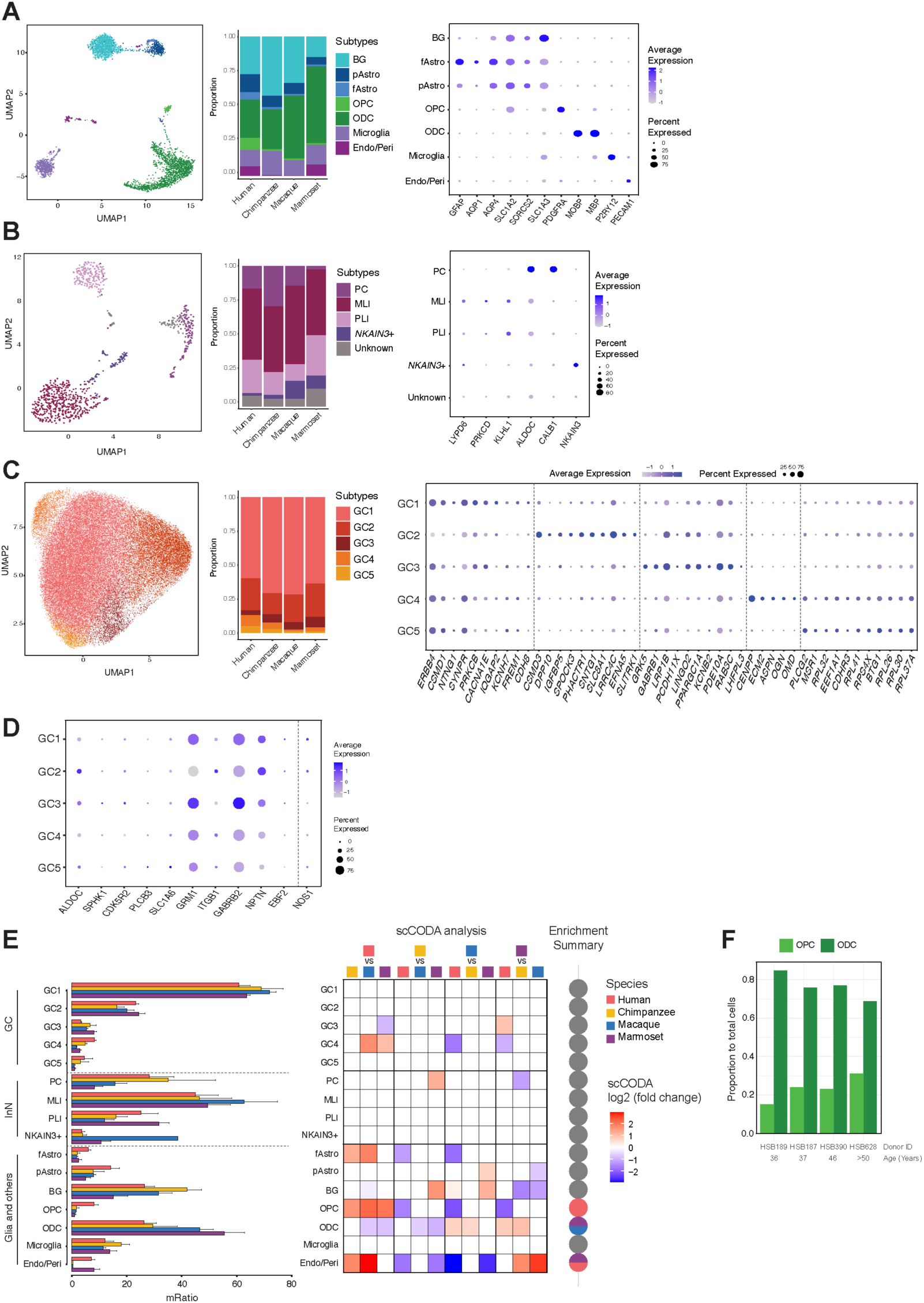
Subclusters of cerebellar cell types and their proportions, related to Figure 1. (A) Proportion of glial cell subclusters and gene expression levels of subcluster markers. pAstro, protoplasmic astrocyte. fAstro, fibrous astrocyte. (B) Proportion of inhibitory neuron subclusters and gene expression levels of subcluster markers. MLI, molecular layer interneuron. PLI, Purkinje layer interneuron. (C) Proportion of GC subclusters and gene expression levels of subcluster markers. (D) Gene expression of cerebellar microzone markers in GC subclusters. (E) Cell type proportions. Left: Bar plots illustrating subtype proportions in each cell type group. To accommodate bias, the proportions were calculated among the following cell type groups: GC, InN, and other non-neural cells. Middle: Heatmap showing scCODA-corrected fold changes of subtype abundance between each pair of species. Non-zero values represent significant changes. Right: Pie charts summarizing species enrichment patterns of subtype abundance. (F) Proportions of OPCs and ODCs across human donors.

**Figure S3.**
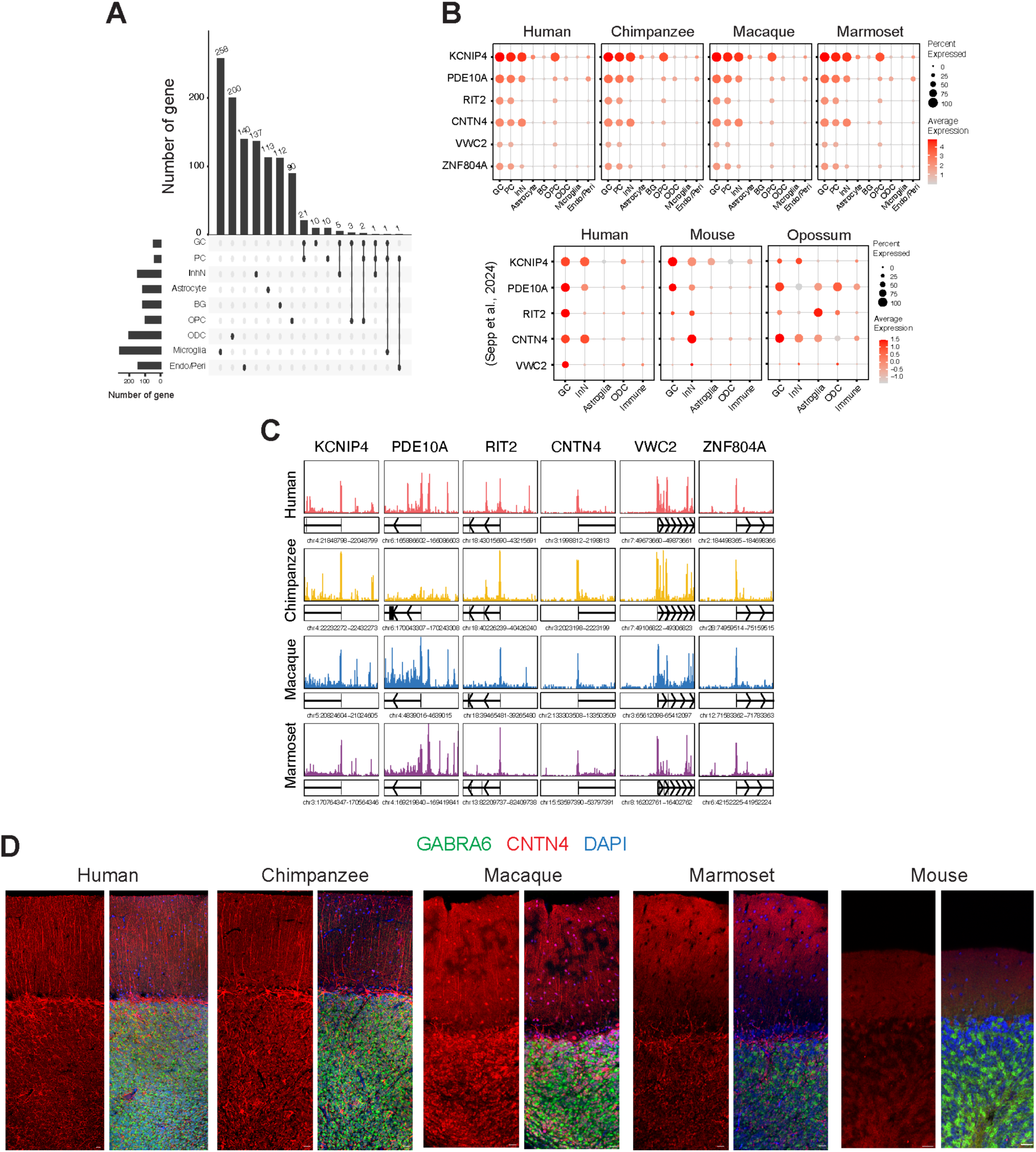
Conserved molecular features of the primate cerebellum, related to Figure 1. (A) Number of genes with conserved expression levels across primates within each cell type. (B) Expression of the six GC-enriched genes that exhibit high expression levels and conserved patterns across primates (Top) and in the adult human, mouse, and opossum cerebellum^17^ (Bottom). (C) Genome browser tracks showing aggregate chromatin accessibility profiles at gene loci of GC-specific conserved expression across primates. (D) Localization of CNTN4 protein in adult cerebellum across four primates and mice.

**Figure S4.**
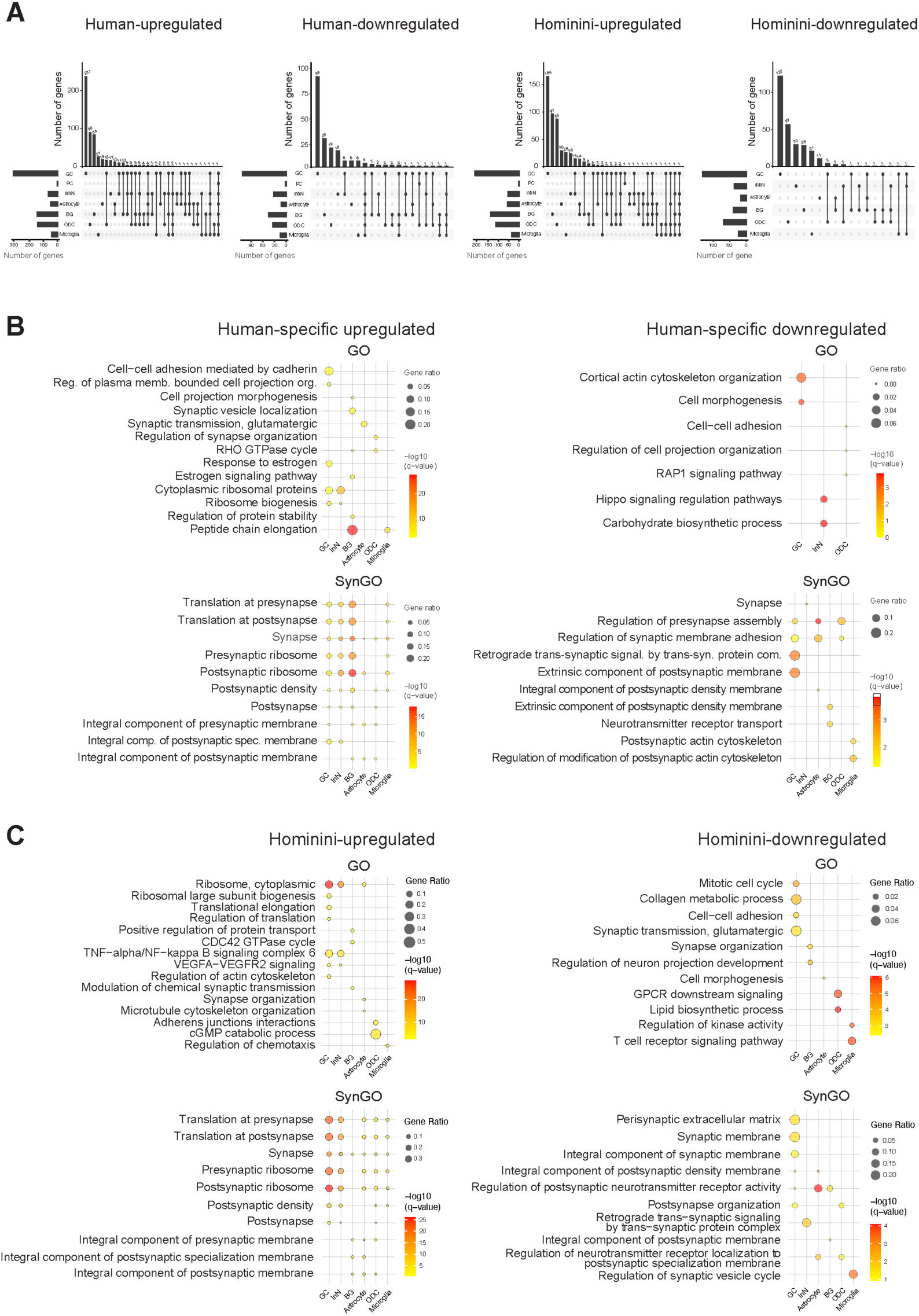
GO and SynGO enrichment of divergent genes, related to Figure 2. (A) Number of genes exhibiting human- and Hominini-specific upregulation and downregulation across cell types. (B-C) Significantly enriched Gene Ontology (GO) and Synaptic GO (SynGO) terms for human-specific upregulated or downregulated genes (B) and for Hominini-specific upregulated or downregulated genes (C).

**Figure S5.**
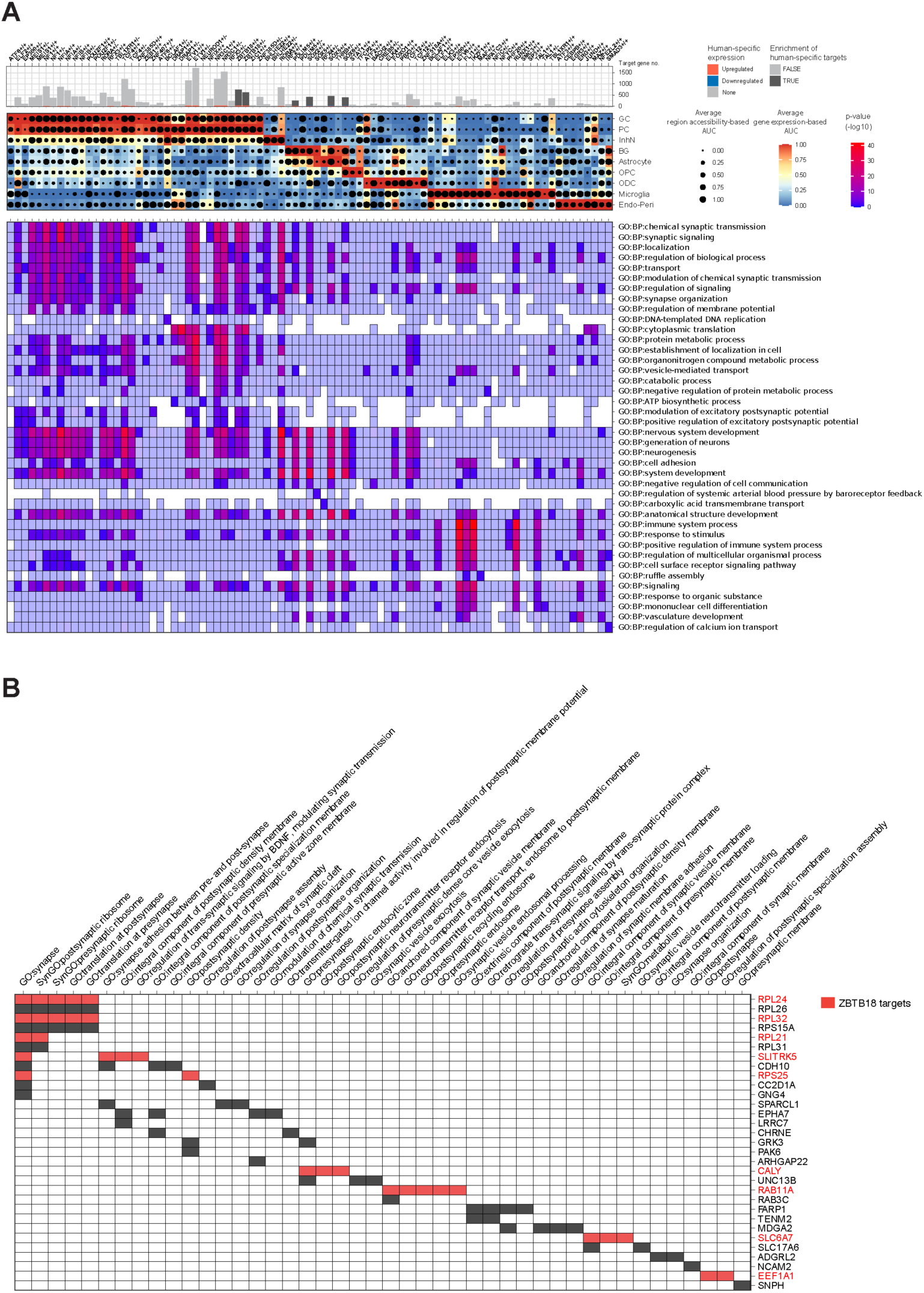
Predicted transcription factor regulatory networks in the human cerebellum, related to Figure 2. (A) Predicted transcription factor (TF) regulatory networks across cerebellar cell types, with Gene Ontology (GO) terms enriched in each regulon. (B) Genes associated with synapse-related GO terms that exhibit human-specific expression. Predicted *ZBTB18* target genes are highlighted in red.

**Figure S6.**
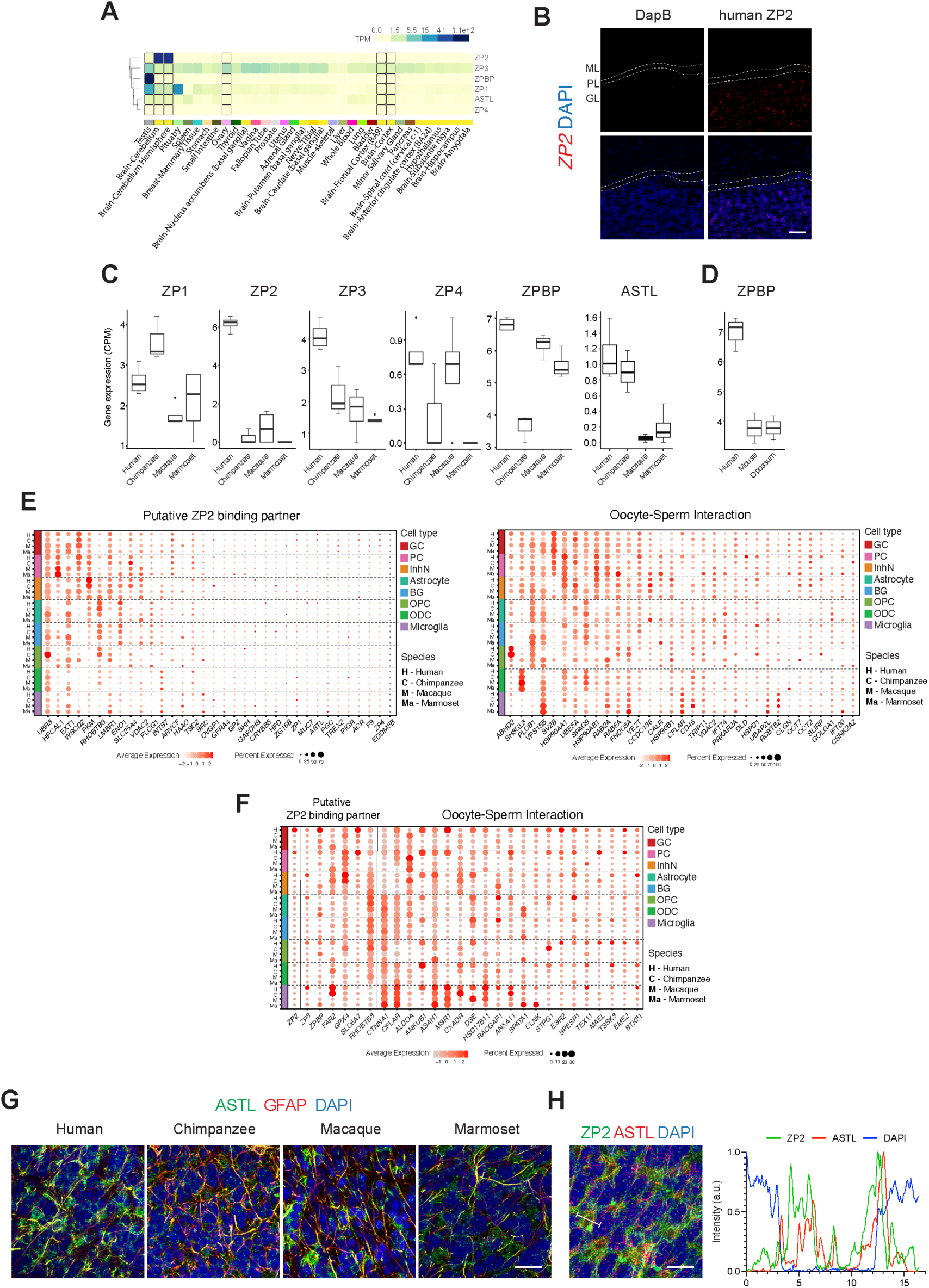
ZP2 and its binding partner expression across species, related to Figure 3. (A) Gene expression of ZP2 family members (*ZP1-4*), *ZPBP*, and *ASTL* across various brain regions and organs in adult humans^33^. (B) ZP2 gene expression in adult human cerebellum by RNAScope in situ hybridization. (C) Gene expression of ZP2 family members (*ZP1-4*), *ZPBP*, and *ASTL* in adult cerebellar GCs across species. (D) *ZPBP* expression in adult cerebellar GCs in humans, opossums, and mice^17^. (E-F) Gene expression of putative ZP2 binding partners and those involved in oocyte-sperm interaction, displaying conserved (E) or divergent (F) expression across cerebellar cell types and species. (G) ASTL and astrocyte marker GFAP expression across species. (H) ZP2 and ASTL expression in the human cerebellum (left) and spatial analysis of their signal intensity along the marked line (right). Scale bars, 20 μm

**Figure S7.**
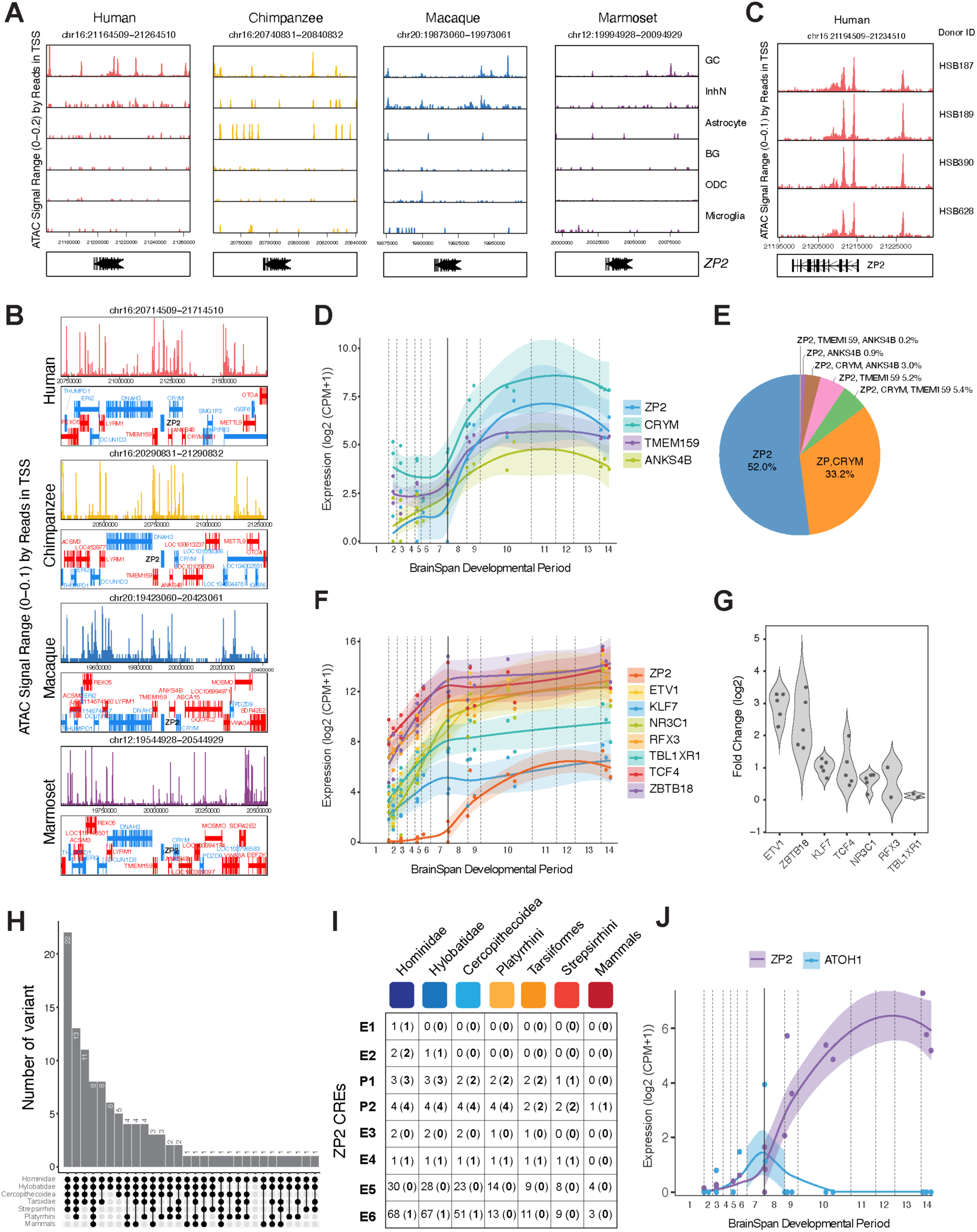
Predicted regulatory networks and cis-regulatory sequence variants underlying human-specific ZP2 expression, related to Figure 4. (A) Cell type–specific chromatin accessibility at the ZP2 locus across primate species. (B) Expanded genomic view of chromatin accessibility surrounding the ZP2 locus in GCs across species. Gene annotations below each track show conserved synteny and gene order. Human GCs exhibit a distinct domain of elevated accessibility spanning TMEM159–ZP2–ANKS4B–CRYM, which is reduced or fragmented in other primates. (C) Donor-level chromatin accessibility at the ZP2 locus in human GCs. (D) Developmental expression of *ZP2*, *CRYM*, *TMEM159*, and *ANKS4B* in GCs in the human cerebellum^17^. (E) Proportion of *ZP2*-positive GCs coexpressing *CRYM*, *TMEM159*, or *ANKS4B* in the adult human cerebellum. (F) Developmental expression of *ZP2* and seven transcription factors predicted to regulate *ZP2* in the human cerebellum^17^. (G) Fold-change expression analysis of the seven transcription factors predicted to regulate *ZP2*, comparing the cerebellum to other brain regions, including the neocortex (NCX), mediodorsal nucleus of the thalamus (MD), hippocampus (HIP), striatum (STR), and amygdala (AMY)^16,48^. *RFX3* and *TBL1XR1* expression in the cerebellum showed differences only when compared to the striatum and hippocampus. (H) Distribution of unfiltered sequence differences present in humans but absent from the indicated mammalian clades. The dataset includes all sites, including those within ≤10 bp of another mutation, located in low-complexity regions, within LTR elements, or associated with INDELs. Bars represent the number of human-specific variants relative to the clades indicated by the connected dots below. A filled dot indicates the absence of the variant in that clade, and connected dots represent clades considered jointly. (I) Cumulative counts of human-specific variants by genomic region and comparative clade. Counts outside parentheses indicate the total number of variants in that region present in humans but absent from the given clade and all intermediate clades along the evolutionary tree. Counts inside parentheses in bold indicate the subset of these variants remaining after applying exclusion criteria: (i) human-specific mutations ≤10 bp apart, (ii) variants in low-confidence areas (e.g., low-complexity or LTR regions), and (iii) variants that are part of INDELs. (J) Developmental expression of *ZP2* and *ATOH1* in GCs during human cerebellar development^17^.

**Figure S8.**
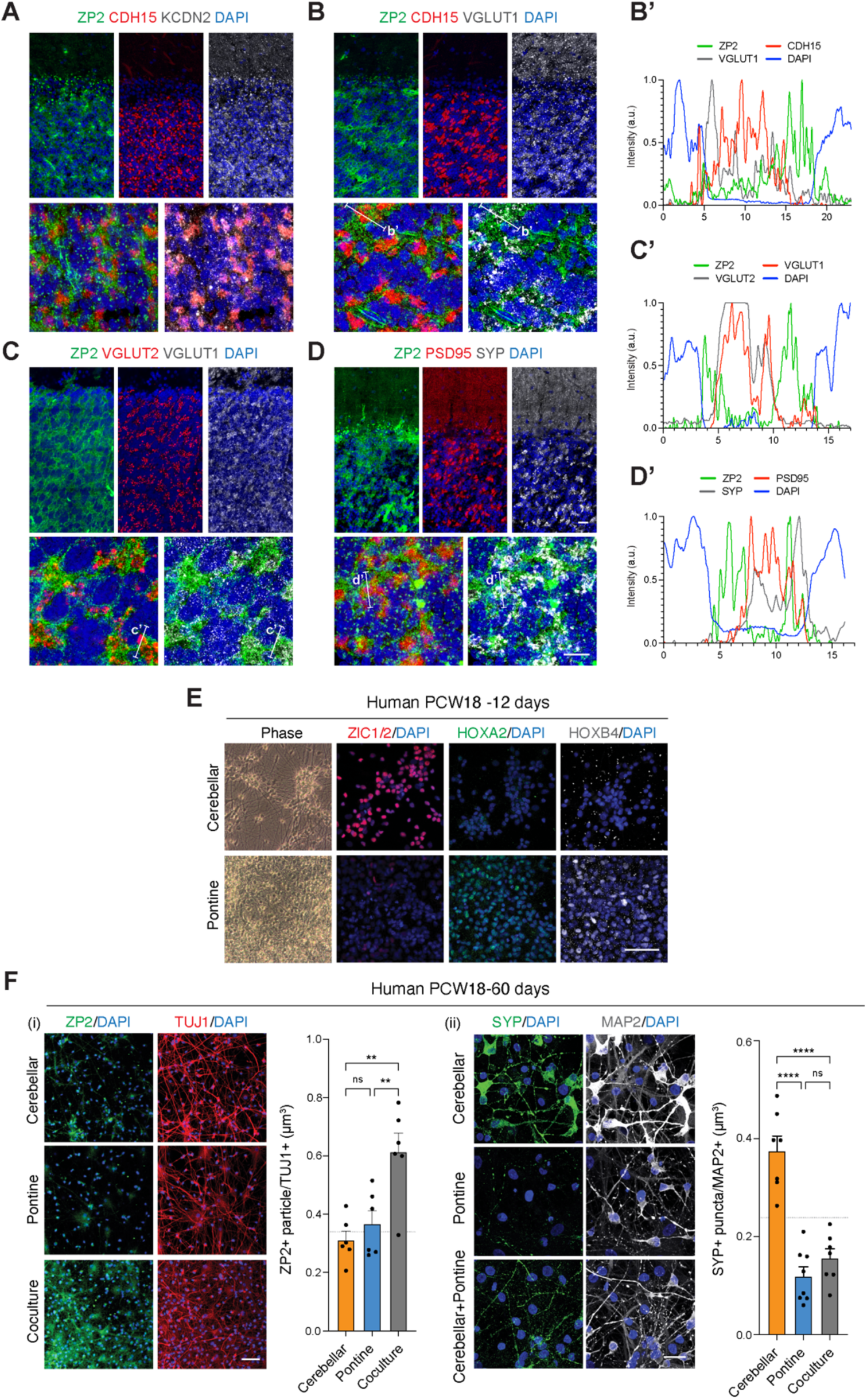
Localization of ZP2 and synaptic proteins in cerebellar glomeruli, related to Figure 5. (A) Immunohistochemical analysis of ZP2 and cerebellar glomerulus maturation markers CDH15 and KCND2. (B) Localization of ZP2, CDH15, and mossy fiber projection marker VGLUT1 at cerebellar glomeruli. Spatial analysis of their signal intensity along the marked line (B’) is shown at the bottom. (C) Localization of dorsal column nuclei and spinocerebellar mossy fiber projection markers VGLUT1 and VGLUT2, along with ZP2, at cerebellar glomeruli. Spatial analysis of their signal intensity along the marked line (C’) is shown at the bottom. (D) Expression patterns of ZP2, presynaptic protein SYP, and postsynaptic protein PSD95 in cerebellar glomeruli. Spatial analysis of their signal intensity along the marked line (D’) is shown at the bottom. (E) Expression of canonical cerebellar markers (ZIC1/2) and pontine markers (HOXA2 and HOXB4) at day 12 in human cerebellar and pontine cell cultures. (F) Human cerebellar, pontine, and cerebello-pontine cell cultures for 60 days in vitro. (i) Induction of ZP2 in cerebello-pontine coculture. (ii) Expression of SYP and MAP2 in human cerebellar and pontine cultures and in cerebello-pontine co-culture. Scale bar, 20 μm. SYP levels exhibit a significant decrease in the co-culture compared to cerebellar cell culture alone. n=7, technical replicates. Significance was assessed using ANOVA for multiple comparisons. ns – not significant, ** p-value < 0.01, **** p-value < 0.0001. Scale bars, (A-D) 20 μm, (E-F) 50 μm.

**Figure S9.**
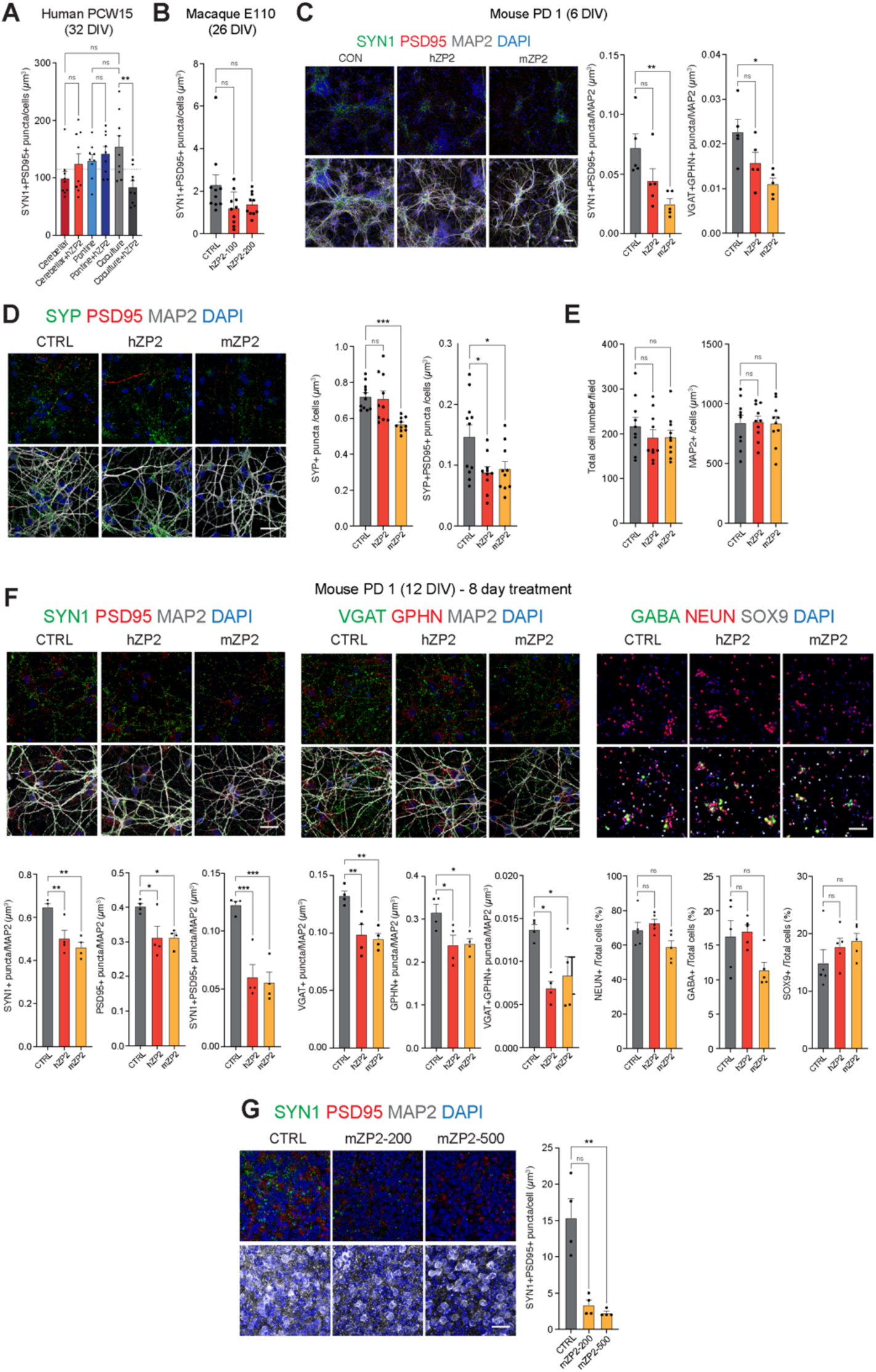
Modulation of synaptic protein expression by ZP2 in cultured cerebellar cells, related to. Figure 6. (A-B) Changes in SYN1-PSD95-positive synaptic puncta in human cerebellar and pontine cell cultures and macaque cerebellar cell cultures following human ZP2 (hZP2) protein treatment. Recombinant hZP2 (100 ng/mL) was added for 48 hours after 32 days of in vitro culture for human cells and 26 days for macaque cells. The total synaptic puncta area (μm^3^) was quantified and normalized by the total number of cells. n=9, multiple fields from 3 technical replicates. (C) Reduced synaptic puncta in mouse cerebellar cells after 48 hours of recombinant mouse (mZP2) and hZP2 treatment. Postnatal day 1 cerebellar cells were cultured for 6 days. Left, Representative images showing SYN1- and PSD95-positive synaptic puncta. Scale bar, 50 μm. Right, Quantification of SYN1-PSD95-positive or VGAT-GPHN-positive synaptic puncta area (μm^3^), normalized to the MAP2-positive area. n=5, independent experiments. (D) Total cell number and MAP2-positive area in mouse cerebellar cells after 48 hours of mZP2 and hZP2 treatment. Postnatal day 1 cerebellar cells were cultured for 6 days. (E) Reduced SYP expression in mouse cerebellar cells after 48 hours of mZP2 and hZP2 treatment. Postnatal day 1 cerebellar cells were cultured for 6 days. (F) Changes of synaptic puncta in mouse cerebellar cells after 8 days of mZP2 and hZP2 treatment. Postnatal day 1 cerebellar cells were cultured for 12 days. n=5, multiple fields from 2 independent experiments. (G) Changes of synaptic puncta in mouse organotypic cerebellar tissue cultures after 48 hours of mZP2 treatment. Cerebellar tissues were collected on postnatal day 21 and cultured for 4 days. CTRL, vehicle (PBS)-treated. n=4, technical replicates. All statistical comparisons performed with Student’s t-test, p-values: ns – not significant, * < 0.05, ** < 0.01, *** < 0.001, **** < 0.0001.

**Figure S10.**
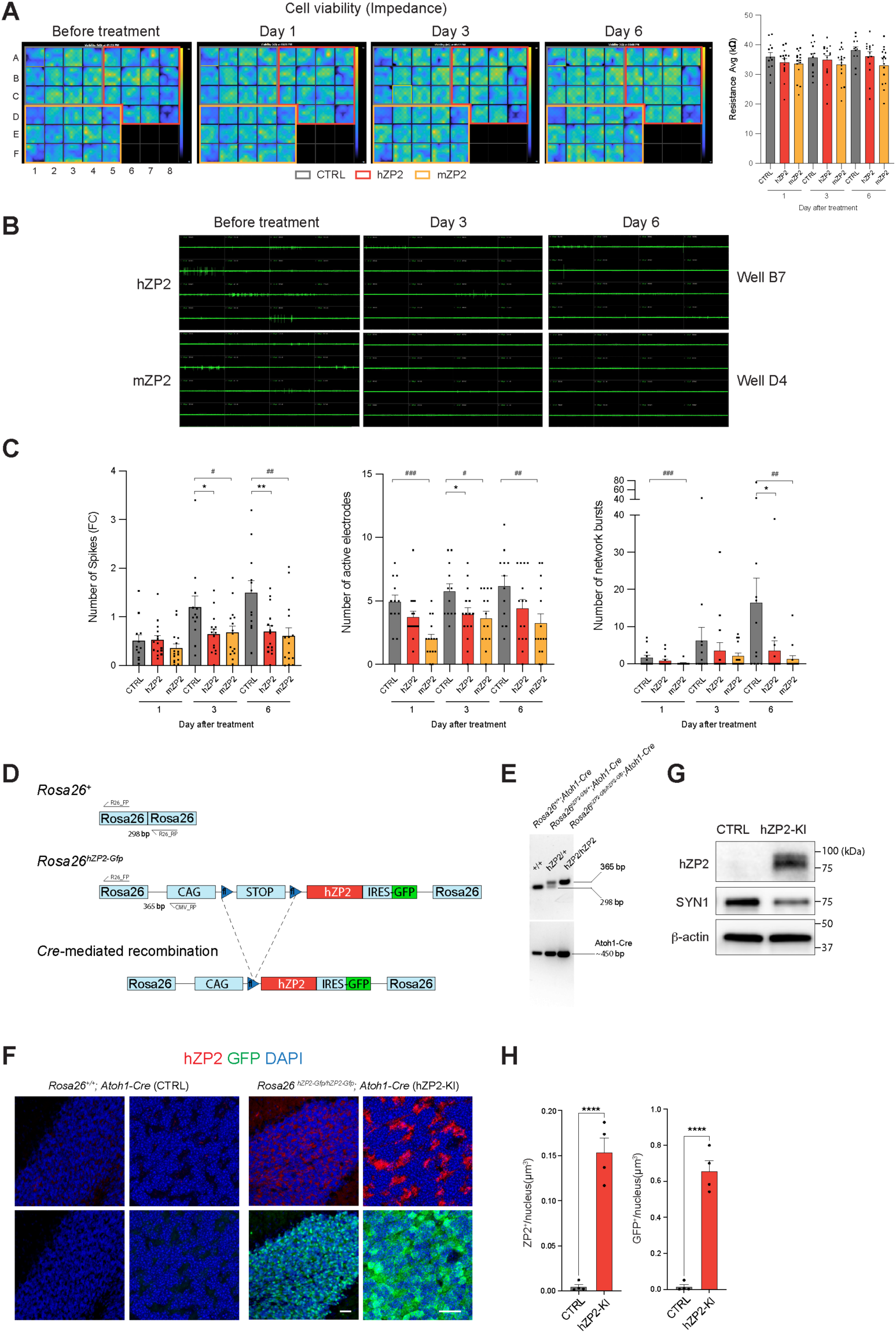
ZP2 suppresses neuronal activity and synapse formation in mouse cerebellar cells in vitro and in the hZP2-KI mouse model, related to Figure 6. (A) Color heatmap visualization of the impedance coverage of the electrodes in the multi-electrode array recording (MEA) plate, showing well-to-well similarity in neuron number and health. Outlines indicate treatment groups. Impedance values are plotted as means in the bar graph. (B) Screenshot views of selected wells display spiking activity for individual electrodes, demonstrating reduced activity following ZP2 treatment. (C) Enhanced display of spiking activity from individual wells (points) and the mean ± SEM (colored bars) from each treatment group. All wells exhibited reduced spiking activity after media change (1-day response). Control wells rapidly recovered to exceed pre-stimulation activity levels, while ZP2-treated wells did not return to pre-stimulation levels even after 6 days. The overall reduction in spiking activity may be due to the silencing of some neurons, as reflected in the number of active electrode plots, which did not show strong statistical significance. The number of network bursts was significantly reduced by treatment on day 6. CTRL, vehicle (PBS)-treated. (D) Schematic showing the Rosa26^+^ wildtype allele (top), the Rosa26^hZP^^2^ conditional knock-in (KI) allele (middle), and the recombined allele after Cre-mediated excision (bottom). The targeted allele contains a CAG promoter, loxP-flanked STOP cassette, hZP2, and IRES-GFP. PCR genotyping regions and expected amplicon sizes (298 bp for control, 365 bp for targeted) are indicated. (E) Genotyping agarose gel showing PCR products used to distinguish Rosa26^+/+^, Rosa26^hZP^^2/+^, and Rosa26^hZP^^2/hZP^^2^ alleles based on 298 bp (control) and 365 bp (targeted) amplicons. A separate PCR detects the presence of the Atoh1-Cre transgene (∼450 bp). (F) Immunostaining of cerebellar sections from control (Rosa26^+/+^; Atoh1-Cre, CTRL) and hZP2-KI (Rosa26^hZP^^2-Gfp/hZP^^2-Gfp^; Atoh1-Cre, hZP2-KI) mice., with GFP marking the hZP2-KI alleles. (G) Western blot showing hZP2 protein expression and reduced SYN1 levels in cerebellar lysates from hZP2 KI mouse compared to control. (H) Quantification of GFP and ZP2 intensity in hZP2-KI and control mice. n=4 mice per group. All statistical comparisons were performed using Student’s t-test, p-values: ns – not significant, CTRL vs. hZP2; *, < 0.05, ** < 0.01. CTRL vs. mZP2; ^#^, < 0.05, ^##^ < 0.01, and ^###^ < 0.001.

**Figure S11.**
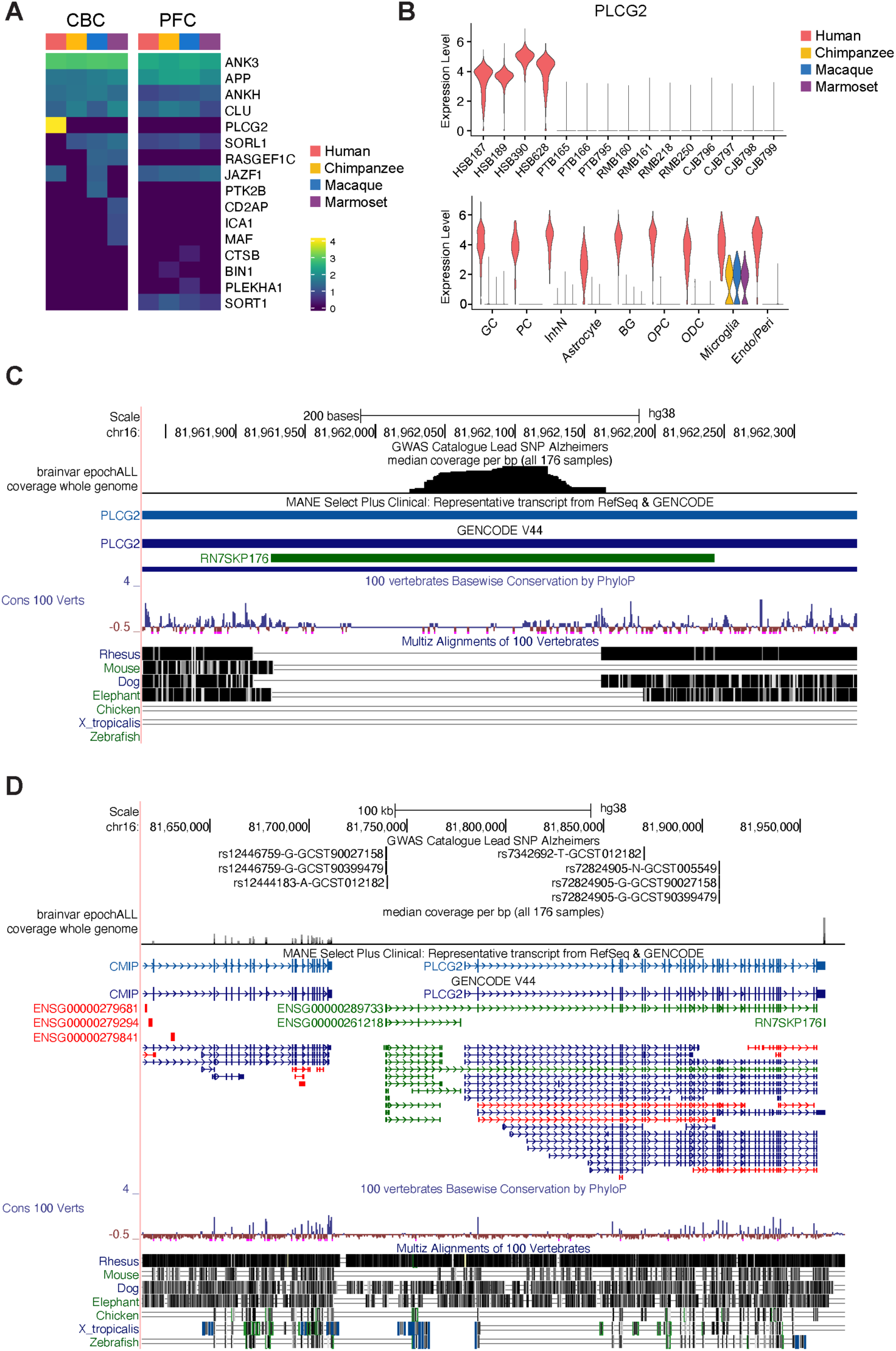
Human-specific signals at *PLCG2* and *RN7SKP176* loci, related to. **Figure 7**. (A) Expression of genes associated with Alzheimer’s disease (AD) in the CBC and PFC across species. (B) Expression of *PLCG2* across donors and cerebellar cell types. (C-D) UCSC Genome Browser views of the GWAS loci in AD around the *PLCG2* gene (C), and expression of this gene and a nearby small RNA pseudogene *RN7SKP176* near the 3’UTR (D).

