## Supplemental Materials and Methods for "Human-specific features of the cerebellum and ZP2-regulated synapse development"

### Human, chimpanzee, rhesus macaque, and marmoset postmortem tissue

Brain samples from humans, chimpanzees, and rhesus macaques were obtained from existing brain banks at the Sestan laboratory. One chimpanzee and four marmoset samples were obtained from the Southwest National Primate Research Center (www.nprc.org), and two chimpanzee samples were obtained from the National Chimpanzee Brain Resource (www.chimpanzeebrain.org). Tissue collection followed guidelines of the Yale Human Investigation Committee (HIC) and NIH ethical guidelines and regulations for the research use of human brain tissue (http://bioethics.od.nih.gov/humantissue.html), as well as the WMA Declaration of Helsinki (http://www.wma.net/en/30publications/10policies/b3/index.html). The use of chimpanzee biomaterials complied with the "NIH Research Involving Chimpanzees (NOT-OD-16-095)."

Tissue specimens were assessed for disease, injury, and gross anatomical alterations. Informed consent was obtained prior to human tissue collection, and non-identifying information was recorded for each specimen. Only samples from subjects without overt clinical pathologies were included.

### Tissue dissections and processing

Samples of the posterolateral cerebellum were dissected from four adult human, three adult chimpanzee, four adult macaque, and four adult marmoset brains (Figure S1, Table S1) and stored at -80 ºC. The collected samples contained all three layers of the cerebellar cortex, without underlying white matter or deep cerebellar nuclei. Where possible, samples from both the right and left cerebellum were pooled. No obvious signs of neuropathological alterations were observed in any of the human and non-human specimens.

### Nuclei isolation

Nuclei were isolated according to published protocols with modifications for cerebellar tissue density[1]. Samples were pulverized in liquid nitrogen using a mortar and pestle (Coorstek #60315, #60317). Then, 10-20 mg of pulverized tissue was added to 3 mL ice-cold lysis buffer (250 mM sucrose, 25 mM KCl, 5 mM $MgCl_2$, 20 mM Tris-HCl, pH 7.5, 0.1%, Igepal CA-630 v/v, 1X protease inhibitors without EDTA, and 1 mM DTT). The suspension was transferred to autoclaved, RNase-free tissue grinders (15 ml volume, Wheaton #357544) on ice and homogenized first with loose, then with tight pestles for 40-50 cycles each.

The suspension was strained through a 70-µm tube top cell strainer (Greiner Bio-One, #542170), pre-wetted with 250 µl of lysis buffer. Next, 3 mL of isolation buffer (25 mM KCl, 5 mM $MgCl_2$, 20 mM Tris-HCL, pH 7.5, 50% Iodixanol v/v, 0.5 % BSA, 0.5% protease inhibitors without EDTA, 0.08 U/µl RNase inhibitors, 1 mM DTT) was added to the filtered homogenate and mixed by inverting ten times. The total suspension was centrifuged at 1000 g in a 15 mL centrifuge tube for 30 min at 4 ºC in a swing-out rotor. Following centrifugation, the supernatant was carefully removed, and the pellet was resuspended in 200 µL of resuspension buffer A (10 mM NaCl, 3 mM $MgCl_2$, 10 mM Tris-HCl, pH 7.5, 0.01% v/v NP40, 0.01% v/v Tween-20, 0.001% v/v Digitonin, 1X protease inhibitors, 1 mM DTT, 1% BSA, 1 U/µL RNase inhibitor). The mixture was pipetted gently 20 times without introducing air. An additional 300 µL of resuspension buffer A was added and mixed gently 20 times with a P1000 pipette without introducing air. The mixture was left on ice, undisturbed, for 5 min. Next, 500 µL of resuspension buffer B (10 mM NaCl, 3 mM $MgCl_2$, 10 mM Tris-HCl, pH 7.5, 0.1% v/v Tween-20, 1 U/µl RNase inhibitor, 1 mM DTT, and 1% BSA) was added to the suspension and pipetted gently 20 times with a P1000 pipette set to 500 µL without introducing air. The mixture was then incubated on ice, undisturbed for 5 minutes.

Nuclei were filtered again with a 70-µm tube top cell strainer, pre-wetted with 20 µL of resuspension buffer B. The filtrate was transferred to 1.5 mL low-binding Axygen tubes, with 30 µL used to determine density by hemocytometer. The remaining filtrate was centrifuged at 500 g for 10 minutes at 4 ºC. The supernatant was carefully removed, and the pellet was resuspended in 1X nuclei buffer supplemented with 1 mM DTT and 1 U/µL RNase inhibitor. Nuclei concentration was adjusted to 4.5 million per mL for nuclei capture.

### Single-nucleus microfluidic capture and library preparation

Nuclei were prepared for capture according to the 10X Genomics Chromium Next GEM Single Cell Multiome ATAC and Gene Expression protocol (PN-1000283). DNA fragments were prepared for ATAC reactions using Multiome ATAC Kit A, following the 10X multiome protocol. Nuclei were processed for the targeted recovery of 10,000 nuclei on the microfluidic Chromium System (10X Genomics), using Multiome Gel Bead Kit A and Multiome GEM Kit A according to the manufacturer's instructions.

Following nuclei capture, samples were placed on ice and transported to the Yale Center for Genome Analysis core facility, where libraries were prepared according to the Chromium Next GEM protocols. Finally, ATAC and RNA libraries were sequenced using paired-end sequencing with single indexing on the HiSeq 4000 platform (Illumina) following the manufacturer's protocols (Illumina, 10X Genomics). To avoid lane bias, multiple uniquely indexed libraries were multiplexed and distributed over several lanes.

**Single-nuclei multiome data processing**

We used CellRanger ARC (v2.0.1) to align the reads to the reference genome for each species. Human reference packages for GRCh38 (GRCh38 Reference-2020-A-2.0.0, May 3, 2021)) were downloaded from the 10x genome webpage (https://support.10xgenomics.com/single-cell-multiome-atac-gex/software/downloads/latest). For chimpanzees, the panTro5 genome sequence file and NCBI gene reference file were downloaded from UCSC (https://hgdownload.soe.ucsc.edu/goldenPath/panTro5/bigZips/). For rhesus macaque, the rheMac10 genome sequence file and NCBI gene reference file were downloaded from UCSC (https://hgdownload.soe.ucsc.edu/goldenPath/rheMac10/bigZips/). For marmoset, the calJac4 genome sequence file and NCBI gene reference file were downloaded from UCSC (https://hgdownload.soe.ucsc.edu/goldenPath/calJac4/bigZips/). For panTro5, rheMac10 and calJac4 genomes, the gene reference gtf files were sorted by gene name and the gene biological type was added based on the Ensembl gene reference file for each species. Then config files were created for the panTro5 genome sequence file and sorted NCBI reference file for CellRanger-ARC mkref. Here, "chrM" was set as non_nuclear_contigs and the input_motifs was set as the same as human GRCh38 reference packages.

We filtered nuclei based on the following criteria: UMI and feature counts more or less than 3 median absolute deviations, more than 5% mitochondrial gene expression, total ATAC fragment count less than 1000, or transcription start site enrichment score less than 4.

**Clustering and cell type annotation**

For the RNA data, raw count matrices were used as input for the Seurat package (v4.0.6)[2,3] for clustering. The raw count matrices were normalized using the Normalize function and scaled using the ScaleData function. Dimensionality reduction and UMAP projection were performed using RunPCA and RunUMAP. The top 21 principal components (PCs) were used UMAP visualization.

Cell types were annotated based on the expression of a curated list of cell type-specific markers (Table S2). We also examined the overlap of top differentially expressed genes across clusters identified by the FindMarkers function and the curated markers. The major cell types include granule cell (*NEUROD1, GABRA6* and *SLC17A7*), Purkinje cell (*ALDOC* and *CALB1*), inhibitory neuron (*GAD1*, *GAD2*, and *SORCS3*), oligodendrocyte (*MOBP*, *MBP*, and *OLIG2*), oligodendrocyte precursor (*PDGFRA* and *PCDH15*), astrocyte (*SLC1A3, SORCS2* and *GFAP*), bergmann glia (*SLC1A3*), microglia (*LRMDA*, *PLXDC2*, and *P2RY12*), endothelial cell (*PECAM1* and *B2M*), and pericyte (*PDGFRB* and *NID1*).

For the ATAC data, open chromatin regions (peaks) were identified using MACS (v2.2.9)[4] for each pseudo-bulk replicate per cell type (q value = 0.05). A consensus set of peaks were generated by iterative merging of peaks from all cell types. Peaks overlapped with blacklist regions in the genome were removed. The peak count matrix was normalized using the log-TF version of the term frequency–inverse document frequency (TF-IDF) transformation, followed by singular value decomposition. The top 30 latent semantic indexing (LSI) components, excluding the first, were used in downstream analysis. The gene activity matrix was created using the GeneActivity function from the ArchR package[5]. The cell type annotation based on RNA data is used for downstream analysis.

**Data integration across species**

For the RNA data, Seurat canonical correlation analysis was used to integrate transcriptome datasets across species. The top 2,000 highly variable genes were identified using the SelectIntegrationFeatures function. Anchors were identified using the FindIntegrationAnchors function with the first 30 PCs. Hierarchical integration of normalized data was accomplished using the IntegrateData function.

For the ATAC data, homologous peaks were identified by reciprocal liftOver between hg38 and the other three reference genomes. The peak matrices for homologous peaks from four species were merged and clustered using LSI reduction as described above.

**Transcriptome correlation with published studies**

To evaluate the concordance of our transcriptomic profiles with public datasets of human and mouse cerebellum[6,7], we performed Spearman's correlation analyses. These analyses were conducted on pseudo-bulk gene expression profiles of the top 2,000 highly variable genes within each dataset. Cell-type transcriptomic signatures from our dataset, which includes cerebellar cells from humans, chimpanzees, macaques, and marmosets, were compared against adult human and mouse cerebellar datasets. For the human public cerebellar dataset, only cells from the adult cerebellar cortex were utilized. For the adult mouse cerebellar dataset, orthologs were retrieved using biomaRt (v2.58.2)[8].

**Subtype abundance comparisons across species**

Cell subtype abundance comparisons across humans, chimpanzees, rhesus macaques, and marmosets were performed using the single-cell Compositional Data (scCODA) analysis algorithm[9]. This model quantified differences in the proportions of cell subtypes within major groups: excitatory GCs, inhibitory neurons, and other non-neural cells. Within these groups, "GC0" for excitatory neurons, "MLI2" for inhibitory neurons, and "Microglia" for non-neural cells were set as the reference subtypes. These cell types demonstrated high presence across samples and minimal variation in their relative abundance, making them robust references for preserving relative abundance changes throughout the analysis. To estimate the posterior distributions of the compositional balances, we implemented Hamiltonian Monte Carlo (HMC) sampling. We set a false discovery rate (FDR) of 0.2 to identify statistically significant differences. For a visual representation of the scCODA results, we employed ggplot2 to create heatmaps that show log2 fold changes in cell subtype proportions between species pairs. Additionally, pie charts were used to visually summarize the relative abundance and distribution of cell subtypes across species, providing an overview of the compositional analysis.

**Identification of species-conserved cell type markers**

For each species, we identified cell type markers through pairwise cell type comparison using the findMarkers function from Seurat (v4.0.6)[2,3] with parameters logfc.threshold=0.25, min.pct=0.3, and max.cells.per.ident=1000. The intersection of differentially expressed genes (DEGs) identified in all the comparisons between the target cell type and other cell types was identified as cell type markers. This approach focuses on identifying cell-type–specific genes that define core cell identity across species. Species-conserved cell type markers were identified as those markers detected in all four species.

**Identification of species-specific genes**

We detected DEGs across pairwise species comparison based on pseudobulk expression profiles using DESeq2 (v1.42.0). This approach reduces cell-to-cell variability and is optimized for robust cross-species comparisons. DEGs were identified as genes with an adjusted p-value less than 0.05 and an absolute log2 fold change greater than 1. Only genes with detectable expression levels were retained, defined as those expressed in more than 2.5% of cells and in more than 20 cells for a given cell type and species.

**Cell type-specific transcriptomic divergence analysis**

To account for the disproportionate abundance of granule cells relative to other populations, equal numbers of each cell type were downsampled for evolutionary comparisons. For each species, 100 cells per cell type were randomly sampled, and divergence was calculated as 1 – Spearman correlation of the average expression of variable features. This sampling was repeated 50 times to generate distributions of divergence values for each combination of cell type and species. Divergence estimates were then summarized as line plots with standard errors.

**Gene ontology (GO) enrichment analysis**

GO enrichment analysis was conducted using FunSet (http://funset.uno/), Metascape (http://metascape.org/gp/index.html, v3.5)[10], and SynGO (https://www.syngoportal.org)[11]. P-values were calculated using the hypergeometric test and adjusted for multiple comparisons using Benjamini-Hochberg correction. Only GO terms with adjusted P-values less than 0.05 were considered significant. Functional enrichment analysis of gene regulatory networks (GRNs) associated with each transcription factor (TF), was performed using g:Profiler (R package gprofiler2_0.2.3)[12], employing the g:SCS multiple testing correction method with a significance threshold of 0.05. To evaluate the enrichment of human-specific genes within these networks, Fisher's exact test was applied, followed by Benjamini-Hochberg correction.

**Identification of cerebellum-specific genes across brain regions**

To identify cerebellum-specific genes across multiple human brain regions, we used the bulk RNA-seq dataset[13], which includes data from the neocortex (NCX), mediodorsal nucleus of the thalamus (MD), hippocampus (HIP), striatum (STR), amygdala (AMY), and cerebellum (CBC). Using DESeq2 (v1.42.0)[14], genes displaying significantly higher expression levels in CBC compared to the other five brain regions were identified as CBC-specific genes (p-value < 0.05). These CBC-specific genes were then ranked based on the minimal log2(FoldChange) among comparisons with the other five regions. The top 50 CBC-specific genes were visualized in Figure 3A.

**Identification of putative peaks linked to target genes**

Putative peaks linked to target genes were identified as either peak-to-peak links (co-accessible peaks to promoter-proximal peak) or peak-to-gene links (peaks whose accessibility correlated with target gene expression). Peak co-accessibility analysis was conducted using Cicero (v1.3.9)[15]. Signac (v1.10.0)[2] was employed to detect peak-to-gene links within 50 kb of the transcription start site. To determine peaks linked to human-specific genes across major cell types, peak-to-peak, and peak-to-gene links within a 50 kb region upstream and downstream of the promoter were calculated for each cell type. Both linked peaks and promoter-proximal peaks were included in downstream analysis. ZP2 cis-regulatory regions were examined for human genomic changes, including human-gained cis-regulatory elements (HgainCREs)[16,17], human ancestor quickly evolved regions (Haqer)[18], human-specific conserved deletions (Hcondel)[19], human accelerated regions (HAR)[20-25], natural selection sites (SelSite)[26-28] and various types of repeat elements obtained from the UCSC genome browser (https://genome.ucsc.edu/cgi-bin/hgTables).

**Generation of gene regulatory networks**

A gene regulatory network (GRN) was constructed from single-nucleus Multiome human data using SCENIC+ (1.0a1)[29], which links transcription factors (TFs) to their putative target genes through regulatory chromosomal regions. The pipeline involves three main steps. First, pycisTopic was used to analyze scATAC-seq data, identifying co-accessible chromatin regions that potentially function as cis-regulatory elements. Next, pycisTarget performed motif enrichment analysis on these regions to associate TFs with their candidate regulatory regions. Finally, SCENIC+ integrated TF expression, chromatin accessibility, and motif information to infer enhancer–gene regulatory networks. For each inferred eRegulon, enrichment scores were calculated at single-cell resolution using the AUCell function from the ctxcore Python package. Specifically, two types of AUC values were computed: (i) regulatory-region accessibility–based AUC, which measures the enrichment of accessible regulatory regions associated with an eRegulon, and (ii) target-gene expression–based AUC, which measures enrichment of its predicted target genes. The resulting per-cell values were averaged across all cells of the same cell type, yielding cell type–level enrichment scores that capture both chromatin accessibility and gene expression contributions. Network visualizations were produced using Cytoscape (3.10.3)[30].

**Multiple sequence alignment and extraction of target regions**

Multiple sequence alignment (MSA) files in MAF format for chromosome 16, containing sequences from 447 mammalian species, were obtained from the UCSC Table Browser[31]. These alignments were generated using the Progressive Cactus aligner[32] and incorporate data from the Zoonomia project and related large-scale comparative genomics studies[33-36]. The mafToRegion tool from kentutils was used to extract aligned sequences corresponding to eight genomic regions of interest.

**Identification of human-specific sequence differences**

Human-specific sequence differences were identified using a custom R script by comparing the human (GRCh38) sequence to those of six great ape species present in the MSA (two Gorilla, two Pongo, and two Pan species). Two categories of differences were defined:(i) Human-specific substitutions: positions where all six great ape species had a nucleotide different from the human base. Only positions in which both the human and all great ape bases were nucleotides (i.e., not alignment gaps) were considered. (ii) Insertions/deletions (indels): (a) Human-specific insertions: nucleotides present in humans but absent in all great apes. (b) Human-specific deletions: gaps in the human sequence where at least one great ape species had nucleotides. Indels were recorded separately from substitution events. This analysis identified 111 human-specific positions (substitutions or indels) across the eight regions.

**Comparative analysis across primate and mammalian clades**

The 111 human-specific positions were further analyzed across all 447 species in the alignment to evaluate their distribution among broader evolutionary clades. Species were classified into the following groups based on taxonomy: Hominidae (n=6), Hylobatidae (n=13), Cercopithecoidea (n=85), Platyrrhini (n=83), Tarsiiformes (n=4),

Strepsirrhini (n=51), and other mammals (n=204). For each position, we determined whether at least one species within each clade shared the human nucleotide. Positions were classified as clade-specific when no species within a given clade matched the human base.

**Transcription factor binding site prediction**

Potential transcription factor binding sites (TFBS) within the eight genomic regions were predicted using FIMO[37] with the non-redundant vertebrate position frequency matrices from the JASPAR 2024 database. Predictions with a p-value $\leq$ 0.01 were retained, and a more stringent q-value $\leq$ 0.01 filter was applied to select high-confidence sites. The resulting TFBS dataset was intersected with the set of human-specific positions identified in the great ape comparison, considering only non-gap positions. Only TFBS containing at least one non-gap human-specific base were retained for downstream analyses.

**Motif disruption analysis**

The potential functional impact of human-specific substitutions on TF binding was assessed using the motifbreakR R package[38]. Only substitution events (no indels) overlapping predicted TFBS were considered. For each site, the human base was computationally replaced with the nucleotide observed in Pan paniscus (bonobo), and the effect on binding affinity was scored using frequency matrices from the MotifDb package and the GRCh38 reference genome. Results were filtered to include only predictions from JASPAR matrices and only TFBS previously identified by FIMO in the same regions. Finally, TFBS were classified according to: (i) whether the binding affinity score for the human reference nucleotide was higher or lower than that for the alternative chimpanzee nucleotide, and (ii) the effect size category assigned by motifbreakR (mild or strong effect).

**Human prenatal post-mortem tissue**

Fresh tissues from prenatal brain specimens (PCW18-HSB755, PCW18-HSB803, PCW21-HSB785) were maintained in ice-cold Hibernate-E (Thermo Fisher Scientific, A1247601) and utilized within 12 to 18 hours post-mortem. Tissues were obtained from the Birth Defects Research Laboratory at the University of Washington (R24HD000836) following parental or next-of-kin consent and approval from the institutional review boards at the University of Washington, Yale University, and the NIH.

**Human and macaque primary cerebellar and pontine cell cultures**

Cerebellar and pontine tissue were dissected and dissociated into single cells for culture. Tissues were mechanically fragmented into small pieces in artificial cerebrospinal fluid (aCSF) supplemented with the following components: 92 mM N-Methyl-D-glucamine (Sigma-Aldrich, M2004), 20 mM HEPES (Sigma-Aldrich, H3375), 5.5 mM glucose (Sigma-Aldrich, G7021), 30 mM sodium bicarbonate (Sigma-Aldrich, S5761), 5 mM sodium L-ascorbate (Sigma-Aldrich, A4034), 2.5 mM potassium chloride (Sigma-Aldrich, P9541), 1.25 mM sodium phosphate (Sigma-Aldrich, S0751), 2 mM thiourea (Sigma-Aldrich, T7875), 3 mM sodium pyruvate (Sigma-Aldrich, P2256), 5.5 mM urea (Sigma-Aldrich, U5128), 10 mM magnesium sulfate (Sigma-Aldrich, M7506), and 0.5 mM calcium chloride (Sigma-Aldrich, 21115). The tissues were then incubated with 2 mg/mL papain (Transnetyx, PAP) for 20-30 minutes at 37 ºC. After enzymatic digestion, 0.1 mg/mL DNase I (Stem Cell Technologies, 07900) was added, and single-cell dissociation was achieved through repeated pipetting with a 1 mL glass pipette. The dissociated cerebellar and pontine cells were plated onto poly-L-ornithine/laminin-coated wells at a density of $2x10^5$ cells per well in 24-well plates (ibidi, 82406), either independently or in co-culture. Cells were cultured with DMEM/F12 (Gibco, 11330-032):Neurobasal (Gibco, 21103-049)(1:1) supplemented with 1X N2 (Gibco, 17502048), 1X B27 (Gibco, 17504044), 1% GlutaMAX (Gibco, 35050061), 1% MEM non-essential amino acid (Gibco, 11140050), 0.1 mM beta-mercaptoethanol (Sigma-Aldrich, M3148), 1 ug/ml heparin (Stem Cell Technology, 7980), 5 ug/ml human insulin (Sigma-Aldrich, I9278), and 1% penicillin-streptomycin (Gibco, 15140122). Cells were cultured for 21 to 60 days. For recombinant human ZP2 experiments, 100 to 200 ng/mL recombinant human ZP2 protein (OriGene, TP319115) was added to the medium, and cells were cultured for 48 hours.

**Mouse primary cerebellar cell cultures**

Primary cerebellar cells were prepared from postnatal day 1 (P1) mouse brains. Pups were decapitated, and their brain were aseptically dissected and placed into ice-cold aCSF. The dissected cerebellum was incubated with 2 mg/mL papain (Transnetyx, PAP) for 20-30 minutes at 37 ºC. Following enzymatic digestion, 0.1 mg/mL DNase I (Stem Cell Technologies, 07900) was added, and the tissue was gently dissociated into single cells by pipetting with a 1 mL glass pipette. Dissociated cerebellar cells were plated onto poly-L-ornithine/laminin-coated wells at a density of $4x10^5$ cells per well in a 24-well plate (ibidi, 82406). Cells were cultured in Neurobasal medium supplemented with

1X B27, 1% GlutaMAX, 30 nM sodium selenite (Sigma-Aldrich, S5261), 5 ug/ml human insulin, 1% penicillin-streptomycin, 10 ng/ml BDNF (PeproTech, 450-02), and 10 ng/ml NT-3 (PeproTech, 450-03). Media were changed every two days. On day 4, cells were treated with 200 ng/mL recombinant human or mouse ZP2 (OriGene, TP510212) for 48 hours.

**Organotypic cerebellar tissue slice culture**

Organotypic cerebellar tissue slices were prepared from a P21 mouse. The pup was euthanized by decapitation, and the brain was aseptically extracted and immediately transferred to ice-cold artificial cerebrospinal fluid (aCSF). The cerebellar tissue was sectioned into 250 µm-thick coronal slices using a vibratome (Leica, VT1200S). The slices were then transferred onto culture inserts (Falcon, 353090) placed in 6-well plates containing 1.5 mL of culture medium per well. The culture medium consisted of Neurobasal-A (Gibco, 108888022) supplemented with 1X B27 (Gibco, 17504044), 1% GlutaMAX (Gibco, 35050061), 5 µg/mL human insulin (Sigma-Aldrich, I9278), 20% horse serum (Gibco, 26050088), and 1% penicillin-streptomycin (Gibco, 15140122). On day 2, the cerebellar slices were treated with mouse ZP2 protein for 48 hours before collection.

**Immunocytochemistry**

Cerebellar and pontine cells were fixed with 4% paraformaldehyde for 10 minutes, followed by three washes with PBS. Cells were permeabilized with 0.3% Triton X-100 (Sigma-Aldrich, T8787) in PBS for 10 minutes at room temperature and blocked with 10% (v/v) normal donkey serum (Jackson ImmunoResearch Laboratories, 017-000-121) in PBS for 1 hour at room temperature. The primary antibody was incubated overnight at 4°C in a blocking solution containing 10% (v/v) normal donkey serum.

Primary antibodies used were: ZP2 (Santa Cruz Biotechnology, sc-390422, 1:200; Thermo Fisher Scientific, PA5-87653, 1:200; Sigma-Aldrich, HPA011296, 1:500), ZIC1/2 (Millipore, ABE1958, 1:400), MAP2 (Novus Biological, NB300-213, 1:2000), MAP2 (Abcam, ab32454, 1:1000), SYN1 (SYSY, 106103, 1:500), PSD95 (Thermo Fisher Scientific, 700902, 1:1000; Cell Signaling Technology, 3450, 1:400), GABA (Sigma-Aldrich, A2052, 1:2000), NEUN (Thermo Fisher Scientific, MA5-33103, 1:500), SOX9 (R&D, AF3075, 1:200), SYP1 (SYSY, 101 006, 1:500), VGAT, (SYSY, 131003, 1:200), GPHN (SYSY, 147 021,1:200), HOXA2 (Abcam, ab222304, 1:200), and HOXB4 (Abcam, ab133521, 1:400).

After three washes with PBS, cells were incubated with secondary antibodies (Alexa Fluor 488-, 594- or 647-conjugated AffiniPure donkey anti-IgG, Jackson ImmunoResearch Laboratories, 1:400) prepared in blocking solution for 2 hours at room temperature. Nuclei were stained with DAPI (1 µg/ml, Sigma-Aldrich, D9542) for 5 minutes at room temperature. Images were captured using a confocal microscope (LSM880, LSM900, Zeiss) and analyzed with z-stack images using Volocity (v.6.3.1) and Spotfire (v.11.2.0) software.

**Intracranial ZP2 protein injection**

Stereotaxis surgeries were performed on P22-P26 CD1 mice (Charles River Laboratories) to inject ZP2 proteins. Preemptive analgesia with Ethiqua (3.25 mg/kg BW) was administered 30 minutes before surgery, followed by anesthesia with a Ketamine-Xylazine mixture (100 mg/kg and 10 mg/kg BW, respectively). The scalp was shaved and cleaned with iodine tincture and ethanol before the mice were placed in a stereotaxic apparatus. Artificial eye ointment was applied to prevent corneal drying. A midline scalp incision was made, and a 0.5 mm drill hole was created in the skull at the cerebellar injection site (coordinates: AP-6.5 mm, ML-1.0 mm, DV-1.0 mm). Using an injection pump system (RWD, R-480) and a glass capillary microneedle, 1.5 µl of human ZP2 protein (170 ng/µl) was injected into the cerebellum at a rate of 15 nl/sec. The microneedle was left in place for 5 minutes post-injection to minimize backflow before being slowly withdrawn. The incision was then closed and sealed with skin glue/tissue adhesive. After 2-4 days, the brains were harvested via transcardiac perfusion with 4% paraformaldehyde. Tissue was sectioned at 70 µm thickness using a vibratome (Leica, VT1000S) for subsequent immunostaining.

**Generation of hZP2-knock-in (KI) mice**

All animal procedures were approved by the Yale University Institutional Animal Care and Use Committee (IACUC) and conducted in accordance with NIH guidelines. C57BL/6J mice of both sexes were group-housed (up to 5 per cage) under controlled environmental conditions (25 °C, 56% humidity, 12-hour light/dark cycle) with ad libitum access to food and water. Animal husbandry was overseen by the Yale Animal Resource Center.

To generate conditional hZP2 KI mice, the coding sequence of the human ZP2 (variant 1) was inserted into the Rosa26 locus under the control of a CAG promoter (composed of a CMV enhancer and chicken β-actin promoter), followed by an IRES-GFP sequence to allow co-expression of hZP2 and GFP from a single transcript. A loxP-flanked

STOP cassette was placed upstream of hZP2 to allow Cre-dependent expression. These Rosa26$^{hZP2\text{-}GFP}$ mice were crossed with Atoh1-Cre mice (Jackson Laboratory, strain #011104) to drive GC lineage-specific expression of hZP2 and GFP. Offspring carrying both Rosa26$^{hZP2\text{-}Gfp}$ and Atoh1-Cre (Rosa26$^{hZP2\text{-}Gfp/hZP2\text{-}Gfp}$; Atoh1-Cre) were used as the experimental group (hZP2-KI), and littermate Atoh1-Cre mice lacking the KI allele (Rosa26$^{+/+}$; Atoh1-Cre) served as controls.

**Immunohistochemistry**

Cerebellum sections (thickness, 50 μm) from human, chimpanzee, macaque, and marmoset brain tissue were mounted on microscope slides. To unmask the antigens, sections were subjected to heat-induced antigen retrieval in R-Buffer A (Electron Microscopy Sciences, 62706-10) using a benchtop antigen retriever (Electron Microscopy Sciences) for 20 minutes. Sections were then allowed to cool to room temperature for 20 minutes and rinsed in PBS. Blocking was done for 1 hour at room temperature in PBS with 5% normal donkey serum (Jackson ImmunoResearch Laboratories, 017-000-121) and 0.5% triton X-100. Incubation with primary antibodies was performed in PBS with 5% normal donkey serum and 0.5% triton X-100 at 4°C overnight.

The following primary antibodies were used at the concentrations indicated: ZP2 (Santa Cruz Biotechnology, sc-390422, 1:200), ZP2 (Sigma-Aldrich, HPA011296, 1:500); CDH15 (Atlas antibodies, HPA009139, 1:500); ZP3 (Novus Biologicals, NBP2-30830, 1:200); VGLUT2 (Cell Signaling, #16066S, 1:500); Synaptophysin1 (Synaptic Systems, 101006, 1:500); ZPBP (Atlas antibodies, HPA058673, 1:200); KCND2 (Almone labs, APC-023-GP, 1:100); VGLUT1 (LSBio, LS-B3839, 1:100); PSD95 (Cell Signaling, 3450S, 1:500); Synapsin1 (Synaptic Systems, 106103, 1:500); GABRA6 (Abcam, ab300069, 1:500); MAP2 (NovusBio, NB300-213, 1:2000), ASTL (antibodies.com, A49810, 1:500), GFAP (SYSY, 173 004, 1:3000), and CALB1 (EnCor Biotechnology, CPCA-Calb, 1:2000).

Secondary antibody incubation was performed in PBS with 5% normal donkey serum and 0.5% triton X-100 for 1 hour at room temperature. Secondary antibodies used were Alexa Fluor 488-, 568-, or 647- conjugated AffiniPure Donkey anti-IgG (1:500; Jackson ImmunoResearch). Nuclei were counterstained with DAPI. The slides were mounted using Vectashield PLUS Antifade mounting medium (H-1900).

To assess spatial intensity distribution, we manually defined regions of interest as areas where cerebellar glomeruli are located between DAPI-positive cells. The fluorescence intensity of each channel was normalized using the maximum value. The images were analyzed using Image J (v.1.54f, http://imagej.nih.gov/ij/).

**MEA recording**

Multielectrode array (MEA) recordings were conducted as previously described[39]. Briefly, the cerebellar cells were dissociated and replated into the wells of an MEA multi-well plate (CytoView MEA 48, M768-tMEA-48W, Axion Biosystems). The spontaneous firing activity of neurons was assessed using a multi-well MEA system (MaestroPro, Axion Biosystems). Each well of the recording plate contained 16 low-noise individually embedded microelectrodes with integrated ground electrodes, forming a 4x4 recording grid of electrodes across a 1x1 mm area. Each electrode had a diameter of 30 μm with 200 μm center-to-center spacing between individual electrodes. The wells of the MEA plate were coated with iMatrix-511 SILK Stem Cell Culture Substrate (Reprocell, NP892-021) for 3 hours at 37°C to facilitate the attachment of the cerebellar cells. A density of $10^5$ cells per well was placed directly over the array of electrodes and allowed to settle before filling with maintenance media. To ensure environmental control during recordings, $CO_2$ concentrations were regulated at 5%, and temperature was maintained at 37°C. A spike detection criterion greater than 6 standard deviations above the background signal was employed to distinguish action potentials from noise.

The effect of recombinant ZP2 protein on the spontaneous activity of cerebellar neurons was assessed before and after treatment. To ensure statistical significance, the individual wells of the multi-well MEA plate were analyzed for total spike counts across all the electrodes within each well. Upon reaching equilibrium in temperature and $CO_2$ levels, a recording was acquired for ten minutes. The wells of the MEA plate were recorded in the ‘pre’ condition, and then the wells were divided into groups for ZP2 treatment.

The plate was removed from the MEA instrument and transferred to a laminar flow hood to apply ZP2 proteins to each group. For the 48-well plates, the maintenance volume of media is 600 μl. ZP2 protein solutions were prepared at three times the final concentrations in the media. Treatment was performed by removing 200 μl of media from each well and then replacing it with 200 μl of the three times concentrated solution, restoring the 600 μl total volume and exposing cells to final ZP2 concentrations of 300 ng/ml.

**Disease gene enrichment measurement**

To measure disease gene expression by cell type, we employed an approach that measures disease gene enrichment as the fraction of disease genes expressed in each cell. This cell-based approach allowed for enrichment measurement at cellular resolution, was self-normalizing, and did not require inputting thresholds or other parameters. We computed this fraction for each cell using the equation:

$GF_i = \frac{D_j}{T_i}$ , where $D_i$ = disease gene UMIs at cell $i$, $T_i$ = total gene UMIs in the same cell, and $GF_j$ was the gene enrichment fraction for the cell

We next aggregated gene enrichment fractions by cell type using the ANOVA test, as implemented in the R 'aov' function, using the gene enrichment fraction as the dependent variable and cell type as the categorical independent variable. The combined enrichment of each cluster was taken to be the ANOVA coefficient estimates as output from `summary.lm`. To identify disease genes in our datasets, we used their Ensembl IDs when present in each disease list to find the correct identification of genes regardless of changes in gene names.

Disease-associated risk gene sets were obtained from OMIM (https://omim.org), SRARI Gene (https://gene.sfari.org), as well as the following sources. ASD and NDD genes were from [40]. Ataxia genes were from the PanelApp "Ataxia and cerebellar anomalies - narrow panel" list (v.4.0), with a GEL status of 3 (https://panelapp.genomicsengland.co.uk/panels/477/). ADHD genes were from [41]. Schizophrenia genes were the "broad fine-mapped set"[42]. Alzheimer's Disease genes were from [43]. MDD genes were from [44]. The gene lists for FTD-ALS, Parkinson's Disease (PD), neuroticism (NEUROT), IQ, bipolar disorder (BD), and multiple sclerosis (MS) were from [45,46].

**Gene enrichment comparison**

To compare clusters' enrichments statistically, we applied a bootstrapping resampling technique. We resampled the cells within each cell type with replacement, recomputing the ANOVA each time for 100 iterations. This process generated a distribution of estimates, from which we derived 95% confidence intervals for each enrichment estimate by cell type.

Each disease gene list contained a different number of genes, leading to a different range of enrichment estimates for each disease. To make these disease enrichments comparable, we normalized the estimates by dividing them by the number of genes for the disease, multiplied by a constant (1000) to bring the estimates closer to 1.

We plotted these enrichment estimates as heatmaps using the ComplexHeatmap package in R. Disease gene sets were split by study type and clustered hierarchically as implemented in ComplexHeatmap. Selected diseases were also shown in a correlation plot for CBC versus PFC datasets as plotted in ggplot2. ComplexHeatmap was also used to plot human-specific genes and other genes found in selected disease sets and disease gene expression by cell type. For the disease gene proportion plots, we defined gene expression conservatively as any gene whose median expression across the dataset was greater than 0.

**QUANTIFICATION AND STATISTICAL ANALYSIS**

Statistical analyses are detailed in the figure legends, with additional information provided in the corresponding Methods details sections. The values for n (number of cells, individuals, experiments, or technical replicates) are listed in Table S1 and the figure legends. Mouse in vitro culture experiments were performed at least three times, with reproducible results in all cases. Data are presented as means ± standard error (SE), with error bars representing SE.
